# Genetically independent phenotype analysis identifies LPA and VCAM1 as drug targets for human ageing

**DOI:** 10.1101/2021.01.22.427837

**Authors:** Paul R. H. J. Timmers, Evgeny S. Tiys, Saori Sakaue, Masato Akiyama, Tuomo T. J. Kiiskinen, Wei Zhou, Shih-Jen Hwang, Chen Yao, The Biobank Japan Project, FinnGen Study, Joris Deelen, Daniel Levy, Andrea Ganna, Yoichiro Kamatani, Yukinori Okada, Peter K. Joshi, James F. Wilson, Yakov A. Tsepilov

**Affiliations:** MRC Human Genetics Unit, MRC Institute of Genetics & Molecular Medicine, University of Edinburgh, Edinburgh, UK; Centre for Global Health Research, Usher Institute, University of Edinburgh, Edinburgh, UK; Laboratory of Theoretical and Applied Functional Genomics, Novosibirsk State University, Novosibirsk, Russia; Laboratory of Glycogenomics, Institute of Cytology and Genetics, Novosibirsk, Russia; Center for Data Sciences, Harvard Medical School, Boston, Massachusetts, USA; Department of Statistical Genetics, Osaka University Graduate School of Medicine, Suita, Japan; Divisions of Genetics and Rheumatology, Department of Medicine, Brigham and Women’s Hospital, Harvard Medical School, Boston, Massachusetts, USA; Program in Medical and Population Genetics, Broad Institute of Harvard and MIT, Cambridge, Massachusetts, USA; Laboratory for Statistical and Translational Genetics, RIKEN Center for Integrative Medical Sciences, Yokohama, Japan; Department of Ophthalmology, Graduate School of Medical Sciences, Kyushu University, Fukuoka, Japan; Institute for Molecular Medicine Finland, University of Helsinki, Helsinki, Finland; Analytic and Translational Genetics Unit, Massachusetts General Hospital, Boston, Massachusetts, USA; Stanley Center for Psychiatric Research, Broad Institute of Harvard and MIT, Cambridge, Massachusetts, USA; Framingham Heart Study, Framingham, Massachusetts, USA; Population Sciences Branch, Division of Intramural Research, National Heart, Lung, and Blood Institute, National Institutes of Health, Bethesda, Maryland, USA; Institute of Medical Science, The University of Tokyo, Tokyo, Japan; Max Planck Institute for Biology of Ageing, Cologne, Germany; Cologne Excellence Cluster on Cellular Stress Responses in Aging-Associated Diseases (CECAD), University of Cologne, Cologne, Germany; Laboratory of Complex Trait Genomics, Department of Computational Biology and Medical Sciences, Graduate School of Frontier Sciences, The University of Tokyo, Tokyo, Japan; Laboratory of Recombination and Segregation Analysis, Institute of Cytology and Genetics, Novosibirsk, Russia

**Keywords:** GWAS, GIP, ageing, frailty, healthspan, lifespan, Mendelian randomisation

## Abstract

The length and quality of life is important to us all, yet identification of promising drug targets for human ageing using genetics has had limited success. Here, we combine six large European-ancestry genome-wide association studies (GWAS) of human ageing traits—healthspan, father and mother lifespan, exceptional longevity, frailty index, and self-rated health—in a principal component framework that maximises their shared genetic architecture. The first principal component (GIP1) is more heritable than the original studies and shows strong genetic correlations with length of life as well as multiple indices of mental and physical wellbeing. We identify 27 genomic regions associated with GIP1, and provide additional, independent evidence for an effect on human ageing for loci near *HTT* and *MAML3* using a study of Finnish and Japanese subject survival. Across the genome, GIP1 associations are enriched in genes involved in haem metabolism and pathways related to transcription, neurogenesis, homeostasis, proteolysis, intracellular signalling, immunity, and the muscle system. Finally, using proteome-wide two-sample Mendelian randomisation and colocalisation, we provide robust evidence for a detrimental effect of blood levels of apolipoprotein(a) (LPA) and vascular cell adhesion molecule 1 (VCAM1) on GIP1. Together, our results demonstrate that combining multiple ageing traits using genetic principal components enhances power to detect biological targets for human ageing.

## Introduction

Ageing affects us all, from the personal, progressive loss of health to the collective burden of chronic age-related disease and frailty on society. In humans, the body undergoes a systemic functional decline after reaching adulthood, which manifests itself as age-related disease, infirmity and eventually death^1^. The factors determining the rate of ageing and death are complex and interlinked, and include genetics, lifestyle, environmental exposures and chance.

Quantifying the ageing process is not straightforward. A variety of ageing-related phenotypes have been studied as proxies, from chronological measurements such as the length of time from birth until occurrence of a major disease (healthspan)^2^ or death (lifespan)^3,4^, to cellular deterioration measurements such as telomere attrition^5^ and loss of Y chromosome^6,7^, to holistic measurements such as the frailty index^8^, encompassing multiple functional impairment indicators^9^. While the genetic component of these ageing-related traits tends to be estimated at less than 15%^2,8,10^, recent progress has been made on characterising this component using large genome-wide association studies (GWAS), and combining similar GWAS to increase statistical power^11^.

The benefit of combining GWAS of several ageing phenotypes, especially in different populations, is the ability to detect biological mechanisms that influence multiple core components of ageing, while downweighing population- and trait-specific features. For example, a recent multivariate analysis of healthspan, parental lifespan, and longevity GWAS found that genetic loci that were not shared between traits often associated with population-specific, behavioural risk factors such as smoking and skin cancer^11^. On the other hand, genetic loci shared between traits were associated with biological pathways such as cellular homeostasis and haem metabolism.

However, to date, large ageing-related trait GWAS have only been combined using MANOVA, which detects genetic variation that is either shared between multiple traits or strongly associated with a single trait^12,13^. This mixture generates heterogeneous SNP effect sizes, complicating the downstream analysis^11^. An alternative is to perform principal component analysis on the genetic covariance between traits and use the component loadings to construct new, genetically independent phenotypes (GIPs)^14^. As their name implies, GIPs capture the genetic covariance between phenotypes while being genetically uncorrelated to each other. In practice, this means the first principal component (GIP1) maximises the genetic overlap between all traits, while each subsequent GIP contains genetic variation distinguishing the traits from each other.

In this study, we cluster 11 large ageing-related trait GWAS by genetic similarity, and explicitly quantify their common and unique genetic architecture using the GIP methodology. We then characterise this common genetic ageing phenotype, identify robust genomic loci, and highlight proteins that may be potential drug targets for improving the length and quality of life.

## Results

### Ageing-related traits cluster based on genetic correlations

We gathered publicly available GWAS summary statistics on ageing-related traits measured in at least 10,000 European-ancestry individuals. These included self-rated health^15^, healthspan^2^, father and mother lifespan^4^, exceptional longevity^16^, frailty index^8^, perceived age^17^, Hannum and Horvath epigenetic age acceleration^18^, telomere length^5^, and mosaic loss of Y chromosome^7^ (**Supplementary Table 1**). A variety of UK and European individuals are represented between these studies, from children (aged 10+) to centenarians (aged 100+), with birth years spanning the 20^th^ and early 21^st^ century. The largest sample consists of UK Biobank participants and their parents (see **Supplementary Note** for details of each GWAS).

Calculating genetic correlations (r_g_) from summary statistics and performing hierarchical clustering based on the magnitude of these correlations, we find the first six traits form a cluster of high genetic similarity (|r_g_| ≥ 0.5; P < 5×10^−15^). In contrast, the epigenetic age acceleration phenotypes correlate with each other (r_g_ = 0.5; 95% CI 0.2–0.8) but show only modest similarity to the traits in the first cluster (|r_g_| ≤ 0.3). The remaining GWAS show low or no correlations with other traits (**Figure 1; Supplementary Data 1**).

**Figure 1:**
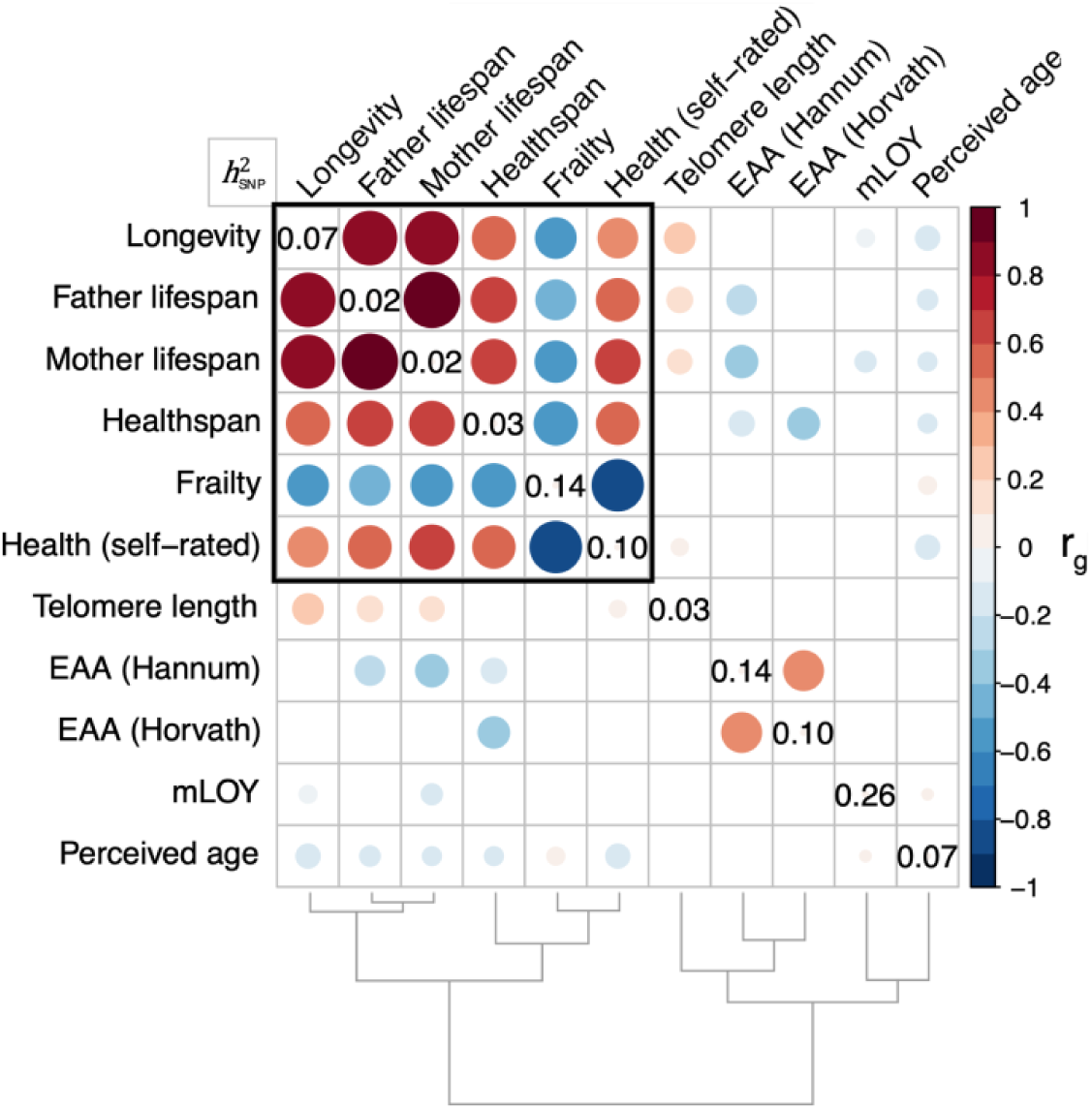
There are strong genetic correlations between measures of the length and quality of life. Diagonal values show the observed-scale SNP heritability of each ageing-related phenotype; off-diagonal circles show the genetic correlation among traits, calculated using High-Definition Likelihood. Blank squares did not pass multiple testing corrections (FDR 5%). The bottom dendrogram shows the hierarchical relationship between traits, based on the magnitude of their genetic correlations. The black box highlights the first cluster of ageing-related phenotypes used in follow-up analyses. EAA—Epigenetic Age Acceleration. mLOY—mosaic Loss of Y chromosome.

Testing the six ageing-related traits in the main cluster for correlations with 728 other traits from the GWAS-MAP platform^19^, we find their genetic similarity may largely be explained through strong, shared genetic correlations (meta |r_g_| ≥ 0.5; P_adj_ < 0.05; P_het_ > 0.05; I^2^ < 50%) with chest pain, cardiovascular disorders, smoking-related disease, type 2 diabetes, and general illness or medication use (**Supplementary Data 2**). However, each core ageing trait also has a number of genetic correlations that differ substantially (see Methods) from the other ageing-related phenotypes and may reflect population or trait-specific risk factors. For example, self-rated health correlates more strongly with physical fitness, body mass index, and a noisy work environment; healthspan correlates more strongly with skin and breast cancers; father lifespan correlates more strongly with hypertension and more weakly with menstrual disorders; mother lifespan uniquely displays a negative correlation with childhood height; longevity uniquely lacks a negative correlation with chronic knee pain; and frailty correlates more strongly with hearing aid usage, daytime napping, and allergic disease (eczema/dermatitis and hayfever/rhinitis). Several ageing-related traits also had uniquely strong correlations with specific medications (**Supplementary Data 2**).

### Genetically independent ageing phenotypes capture distinct elements of wellbeing

We combined the six GWAS in the main correlation cluster using the loadings from the principal components of the genetic correlation matrix (**Supplementary Figure 1**), yielding association summary statistics for six GIPs (available at https://doi.org/10.7488/ds/2972). As expected, GIP1 is the most heritable of the GIPs (h^2^_SNP_ = 0.20; SE = 0.005), capturing over 70% of the genetics of healthspan, parental lifespan, and longevity, and 90% of the genetics of self-rated health and the inverse of frailty (henceforth referred to as “resilience”) (**Figure 2**). A leave-one-out analysis confirms that the GIP analysis is highly robust to the selection of GWAS, with the genetic architecture of GIP1 remaining largely the same after excluding any one of the core ageing-related trait GWAS (range r_g_ GIP1 with GIP1-resilience = 0.950 [SE 0.024] to r_g_ GIP1 with GIP1-healthspan = 0.996 [0.023]) (**Supplementary Table 2**).

**Figure 2:**
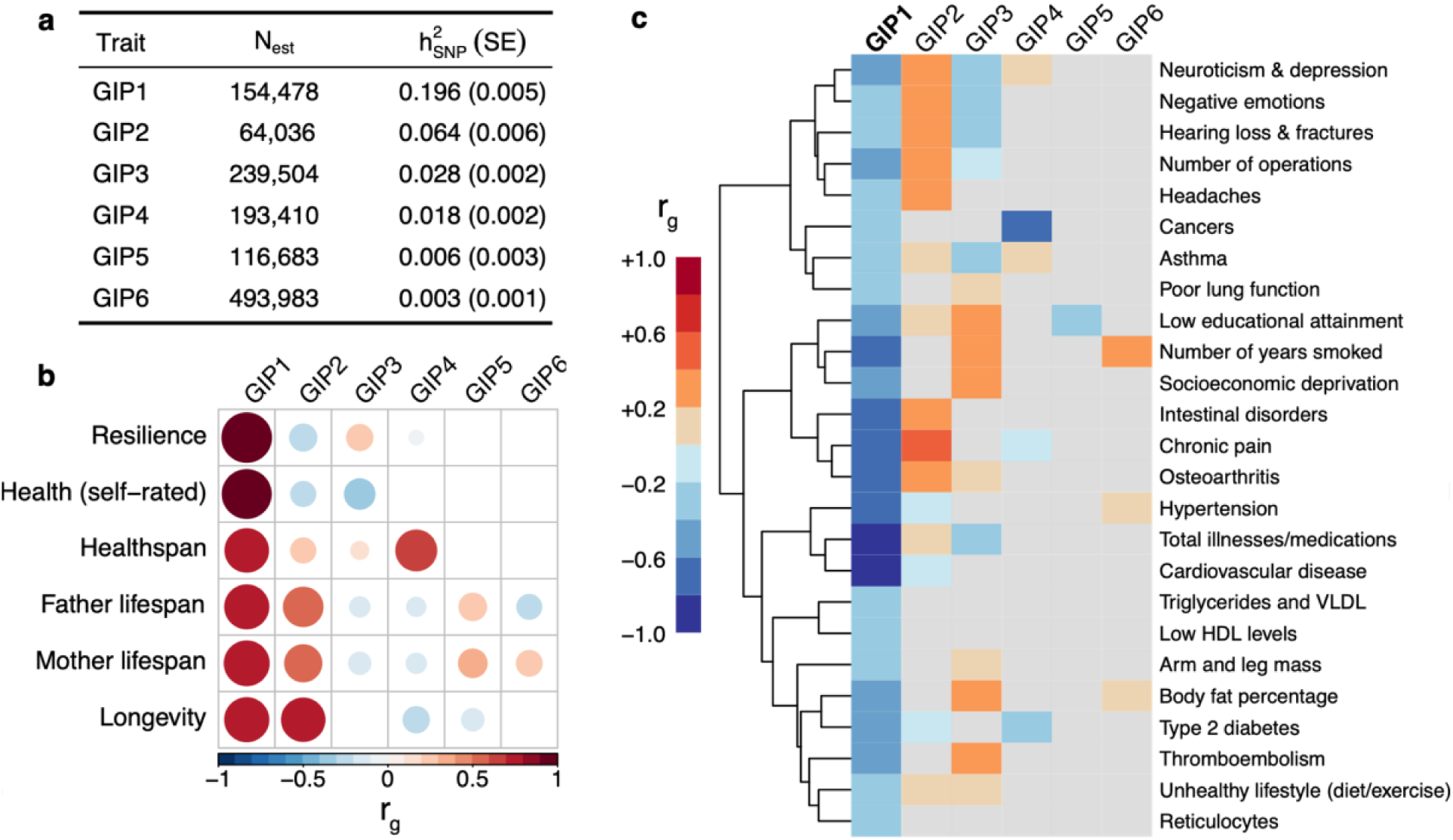
Phenome-wide genetic correlations show GIP1 captures both physical and mental wellbeing. a) Table showing the estimated sample size (N_est_) and SNP heritability (h^2^_SNP_) of each GIP GWAS, with standard errors in parentheses. b) Genetic correlations between GIP GWAS and the core ageing-related trait GWAS used to construct them. Blank values failed to pass nominal significance (P ≥ 0.05). c) GWAS of 402 phenotypes measured in European-ancestry individuals from the GWAS-MAP platform have strong and significant associations with GIP1 (P < 1.5×10^−5^ and |r_g_| > 0.25). These 402 traits are summarised here in 25 groups, based on hierarchical clustering of the magnitude of pairwise genetic correlations. Each group is manually annotated with a label describing the traits within the cluster, and displays the values of the most informative trait (i.e. the trait with the highest total Z score across GIPs). Values failing to pass nominal significance (P ≥ 0.05) are greyed out. The dendrogram displayed here shows the hierarchical relationship between the most informative trait in each cluster. See **Supplementary Data 3** for the full list of correlations.

Apart from the genetic correlations with its component traits, GIP1 shows significant correlations with 459 out of 729 traits tested (see Methods). These suggest that, in addition to length of life, GIP1 also captures mental and physical wellbeing. For example, among the largest negative GIP1 correlations are mental illness, taking medications, and diseases of old age (such as cardiometabolic disorders, cancers, and osteoarthritis) (r_g_ ≤ –0.5; P_adj_ < 1×10^−9^). Inversely, fitness and education are among the largest positive correlations (r_g_ ≥ 0.5; P_adj_ < 1×10^−10^). GIP1 also shows moderate negative correlations with infectious diseases, including N39 (International Classification of Diseases 10th Revision) urinary tract infections (r_g_ = – 0.66; 95% CI –0.42 to –0.90; P_adj_ = 3×10^−4^), coughing on most days (r_g_ = –0.39; –0.30 to –0.49; P_adj_ = 1×10^−11^), and severe COVID-19 hospitalisation (i.e. resulting in respiratory support or death) (r_g_ = –0.33; –0.20 to –0.46; P_adj_ = 0.004). However, GIP1 also retains some strong correlations with socioeconomic factors, such as smoking behaviours (e.g. current tobacco smoking r_g_ = –0.50; –0.45 to –0.55; P_adj_ = 4×10^−92^) and having a job involving manual or physical work (r_g_ = –0.49; –0.44 to –0.54; P_adj_ = 2×10^−89^) (**Figure 2; Supplementary Data 3**).

The remaining GIPs have much lower heritability (h^2^_SNP_ < 0.065) and are of less interest to this study as they capture genetic variance shared between fewer traits and do so with higher degrees of uncertainty. In short, GIP2 correlates mainly with non-lethal causes of poor self-rated health, such as chronic pain (neck, back, stomach), negative emotions, and hearing problems (59 traits, P_adj_ < 0.05); GIP3 correlates mainly with measures of socioeconomic deprivation and body composition (73 traits); and GIP4 captures healthspan-specific correlations with cancer and diabetes (7 traits). While GIP5 and GIP6 are underpowered due to their low heritabilities, they appear to distinguish parental lifespan from longevity (possibly through educational attainment), and father lifespan from mother lifespan (possibly through risk taking and cardiovascular factors), respectively (**Figure 2**).

### Characterising the genomics of lifelong mental and physical health

Across the genome, 27 loci pass the Bonferroni-adjusted genome-wide significance threshold (P < 5×10^−8^/6) in the GIP1 GWAS. The strongest lead SNPs in these loci are rs429358 (P = 2×10^−40^) and rs660895 (P = 2×10^−22^), located nearest to *APOE* and *HLA-DRB1/DQA1*, respectively (**Figure 3; Table 1**). The majority of lead SNPs are strongly associated with self-rated health and resilience, in line with the large loadings of these traits in the construction of GIP1 (**Figure 3; Supplementary Table 3)**. Loci near *APOE, HLA-DRB1/DQA1, LPA*, and *CDKN2B/-AS1* have previously been validated using the same trait in an external cohort^2–4^. For the remaining 23 loci, we measured lead SNP effects on FinnGen participant survival (Release 5; N = 203,244; 6.94% deceased) and BioBank Japan participant survival (N = 135,983; 24.1% deceased), to provide additional evidence of their association with human ageing traits in independent samples. Combining both cohorts to achieve adequate power, we find GIP1-increasing alleles of lead SNPs near *HTT* and *MAML3* have a protective effect on survival in these cohorts (one-sided P_adj_ < 0.05), increasing average lifespan by around 2.53 (95% CI 0.91 to 4.15) and 2.51 (0.79 to 4.23) months per allele, respectively. While we are underpowered to confirm the remaining 21 loci individually, we find that collectively, their GIP1-increasing alleles are also associated with increased Finnish and Japanese survival (one-sided P < 0.05) (**Supplementary Table 4**).

**Figure 3:**
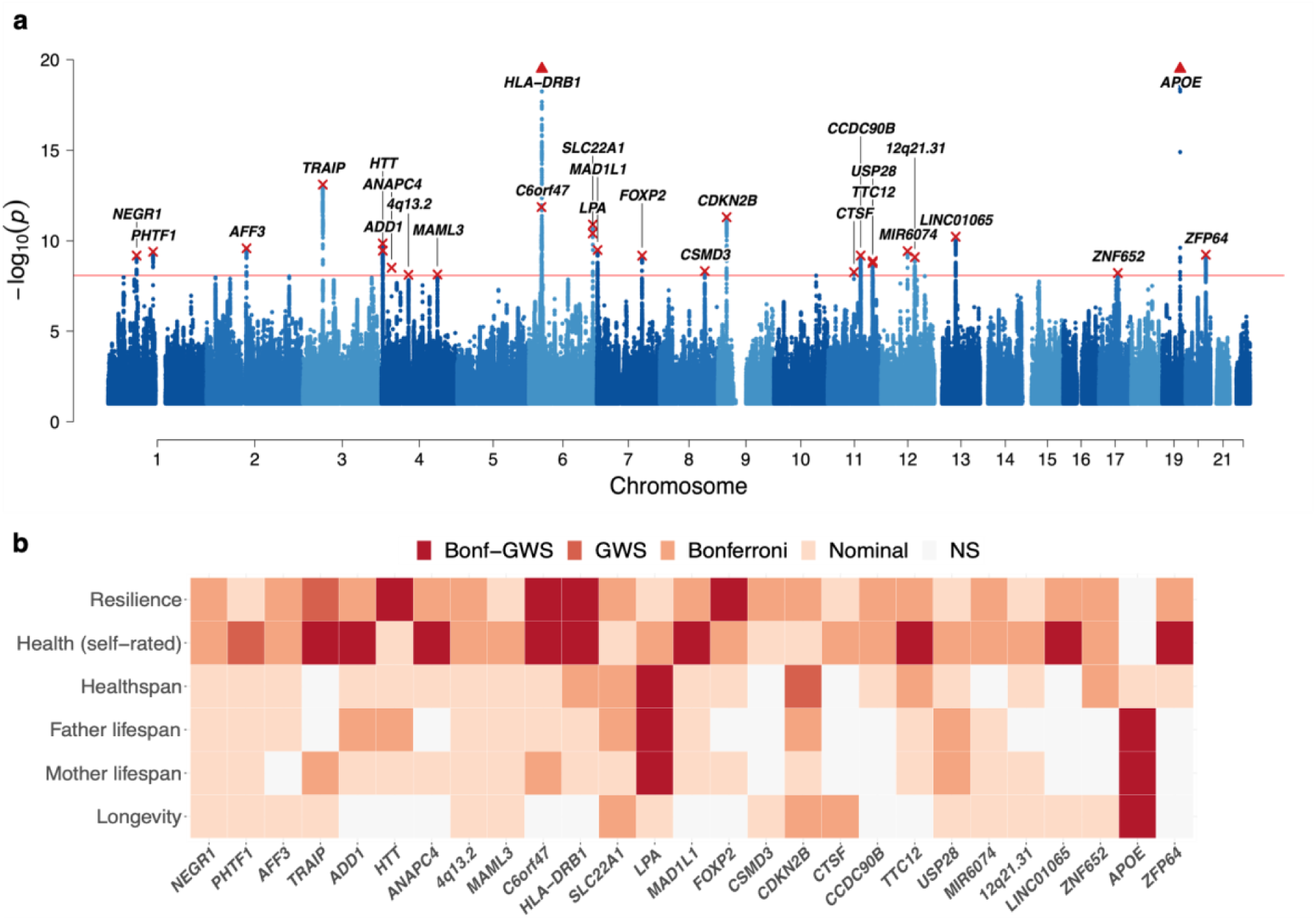
Twenty-seven independent genomic loci are associated with the shared genetic component of ageing-related GWAS. a) Manhattan plot showing associations of SNPs across the genome with GIP1, with the y-axis showing the strength of the association and the x-axis the genomic position of the SNP. The red line represents the Bonferroni-adjusted genome-wide significance threshold of 5×10^−8^/6. The lead SNP of each independent locus is marked with a red cross and annotated with the nearest (upstream) gene, or the cytogenetic band if there are no genes within 250 kb. The y-axis has been capped at P = 1×10^−20^. Loci which exceed this cap and are represented as triangles: *APOE* (P = 1.5×10^−40^) and *HLA-DRB1* (P = 1.4×10^−22^). b) Significance of the association of each lead SNP with the ageing-related trait GWAS used to construct GIP1. Bonf-GWS: Bonferroni-adjusted genome-wide significant (P < 5×10^−8^/6). GWS: Genome-wide significant (P < 5×10^−8^). Bonferroni: P < 0.05 adjusted for 6 traits and 27 SNPs (P < 3×10^−4^). Nominal: P < 0.05. NS: Not significant.

**Table 1:** Twenty-seven independent genomic loci are associated with the first genetic principal component of ageing-related trait GWAS (GIP1). Loci were defined as 500 kb regions centred on a lead genome-wide significant SNP (P < 5×10^−8^/6) in linkage equilibrium (r^2^_EUR_ < 0.1) with other lead locus SNPs. Nearest gene(s)— Closest genes upstream/downstream to the lead SNP (within 250 kb), or if none, the closest cytogenetic band. rsID—The lead SNP within the locus. Chr—Chromosome. Position—Base-pair position (GRCh37). A1—Effect allele, associated with higher GIP1. Freq1—Allele frequency of the effect allele in UK Biobank. β1—Effect estimate (and standard error) of the A1 allele on GIP1 in standard deviation units. P—Nominal Wald-ratio P value. Het—Evidence of heterogeneity: asterisks indicate GIP1 effect size changes significantly when leaving out one of the core ageing traits from GIP1 calculation (* all but one leave-one-out effects are the same, ** all but two effects are the same, *** three or fewer effects are the same).

| Nearest gene(s) | rsID | Chr | Position | A1 | Freq1 | β1 (SE) | P | Het |
| --- | --- | --- | --- | --- | --- | --- | --- | --- |
| *NEGR1* | rs2815748 | 1 | 72816147 | G | 0.20 | 0.028 (0.004) | $7 \times 10^{-10}$ | |
| *PHTF1* | rs1230682 | 1 | 114293526 | A | 0.63 | 0.023 (0.004) | $4 \times 10^{-10}$ | |
| *AFF3* | rs7609078 | 2 | 100490363 | A | 0.37 | 0.023 (0.004) | $3 \times 10^{-10}$ | |
| *TRAIP* | rs2271961 | 3 | 49878113 | T | 0.50 | 0.027 (0.004) | $8 \times 10^{-14}$ | |
| *ADD1* | rs16843603 | 4 | 2928577 | C | 0.28 | 0.025 (0.004) | $1 \times 10^{-10}$ | |
| *HTT* | rs362273 | 4 | 3227419 | G | 0.32 | 0.024 (0.004) | $4 \times 10^{-10}$ | |
| *ANAPC4* | rs34811474 | 4 | 25408838 | A | 0.23 | 0.025 (0.004) | $3 \times 10^{-9}$ | |
| 4q13.2 | rs10434248 | 4 | 67842921 | A | 0.57 | 0.021 (0.004) | $8 \times 10^{-9}$ | |
| *MAML3* | rs56172573 | 4 | 140919381 | C | 0.38 | 0.021 (0.004) | $7 \times 10^{-9}$ | |
| *C6orf47/GPANK1* | rs805262 | 6 | 31628733 | C | 0.52 | 0.027 (0.004) | $1 \times 10^{-12}$ | |
| *HLA-DRB1/DQA1* | rs660895 | 6 | 32577380 | A | 0.79 | 0.043 (0.004) | $1 \times 10^{-22}$ | * |
| *SLC22A1/A2* | rs9456508 | 6 | 160598596 | T | 0.98 | 0.082 (0.012) | $4 \times 10^{-11}$ | ** |
| *LPA* | rs118039278 | 6 | 160985526 | G | 0.92 | 0.045 (0.007) | $1 \times 10^{-11}$ | *** |
| *MAD1L1/SNORA114* | rs11764780 | 7 | 2020904 | C | 0.19 | 0.029 (0.005) | $3 \times 10^{-10}$ | |
| *FOXP2* | rs12705966 | 7 | 114248851 | G | 0.34 | 0.023 (0.004) | $7 \times 10^{-10}$ | |
| *CSMD3* | rs560719 | 8 | 113032374 | T | 0.50 | 0.021 (0.004) | $5 \times 10^{-9}$ | |
| *CDKN2B/-AS1* | rs9632885 | 9 | 22072638 | G | 0.52 | 0.025 (0.004) | $5 \times 10^{-12}$ | ** |
| *CTSF* | rs2924807 | 11 | 66341005 | G | 0.50 | 0.021 (0.004) | $5 \times 10^{-9}$ | |
| *CCDC90B/DLG2* | rs2512690 | 11 | 83145469 | C | 0.34 | 0.023 (0.004) | $7 \times 10^{-10}$ | |
| *TTC12/ANKK1* | rs2186800 | 11 | 113242860 | A | 0.54 | 0.022 (0.004) | $2 \times 10^{-9}$ | |
| *USP28/HTR3B* | rs61907878 | 11 | 113751052 | C | 0.90 | 0.037 (0.006) | $1 \times 10^{-9}$ | |
| *MIR6074* | rs7306710 | 12 | 66376091 | T | 0.48 | 0.023 (0.004) | $4 \times 10^{-10}$ | |
| 12q21.31 | rs6539846 | 12 | 84811217 | A | 0.49 | 0.022 (0.004) | $8 \times 10^{-10}$ | |
| *LINC01065* | rs8002970 | 13 | 53924489 | C | 0.45 | 0.024 (0.004) | $6 \times 10^{-11}$ | |
| *ZNF652/PHB* | rs28394864 | 17 | 47450775 | G | 0.54 | 0.021 (0.004) | $6 \times 10^{-9}$ | |
| *APOE* | rs429358 | 19 | 45411941 | T | 0.84 | 0.066 (0.005) | $2 \times 10^{-40}$ | *** |
| *ZFP64* | rs67442863 | 20 | 51031131 | T | 0.18 | 0.029 (0.005) | $6 \times 10^{-10}$ | |

Again, we find GIP1 genetics are highly stable: when performing a leave-one-out analysis of the core ageing traits, the lead SNP effect of most GIP1 loci does not change significantly. The exceptions are for previously replicated loci and *SLC22A1/A2*, where exclusion of one of the traits can reduce the GIP1 effect size, although loci near *HLA-DRB1/DQA1, LPA*, and *CDKN2B/-AS1* remain genome-wide significant in all leave-one-out scenarios (**Supplementary Data 4**).

We further looked up all lead GIP1 SNPs and close proxies (r^2^_EUR_ ≥ 0.8) in PhenoScanner^20^ and the GWAS catalog^21^, excluding associations discovered solely in UK Biobank, which showed 24 out of 27 GIP1 loci had previously been associated with one or more traits at genome-wide significance. The majority of these loci were associated with cardiometabolic, immune-related, or neuropsychiatric disorders, although several were also associated with measures of educational attainment and household income. Of specific interest are loci near *APOE, HLA-DRB1/DQA1, CDKN2B/-AS1*, and *ZNF652/PHB*, which show GIP1-increasing alleles are associated with a reduction in multiple diseases but do not appear to associate with socioeconomic measures, suggesting these loci largely capture intrinsic sources of ageing (**Supplementary Data 5**).

Aggregating SNP association statistics across the genome into gene scores using the Pathway Scoring Algorithm (PASCAL)^22^, we find significant enrichment in the scores of 1,426 genes (FDR 5%) (**Supplementary Data 6**). High scoring genes appear overrepresented in the haem metabolism hallmark gene set, as well as 432 gene ontology pathways (FDR 5%). These gene ontology pathways cluster into 29 groups, related to transcriptional regulation; neuronal development, organisation, and function; chemical homeostasis; cellular growth, differentiation, and apoptosis; proteolysis; intracellular signalling and transport; immune system development; the muscle system and its response to inactivity; and lipoprotein metabolism (**Supplementary Data 7**). Similarly, GIP1 heritability appears to be enriched in genomic regions containing histone marks associated with the central nervous system and adrenal/pancreatic tissues (**Supplementary Table 5**).

### Causal inference of blood protein levels on healthy life

Next, we used a set of well-validated blood protein quantitative trait loci (pQTL) for 857 proteins as genetic instruments in a two-sample Mendelian randomisation and colocalisation framework to infer putative causal links between protein levels and GIP1 (**Supplementary Data 8)**. We find robust evidence (see Methods) for a detrimental effect of the levels of four proteins in blood on GIP1 (**Table 2**). Three of these proteins—Apolipoprotein(a) (LPA), Olfactomedin-1 (OLFM1), and LDL Receptor Related Protein 12 (LRP12)—were instrumented by a cis-pQTL and were encoded by genes that appeared significantly enriched in the gene score analysis. The remaining protein, Vascular Cell Adhesion Molecule 1 (VCAM1), was instrumented by a trans-pQTL shared with Beta-2-microglobulin (B2M); however, only VCAM1 colocalised with the signal at this locus (**Supplementary Data 8**).

**Table 2:**
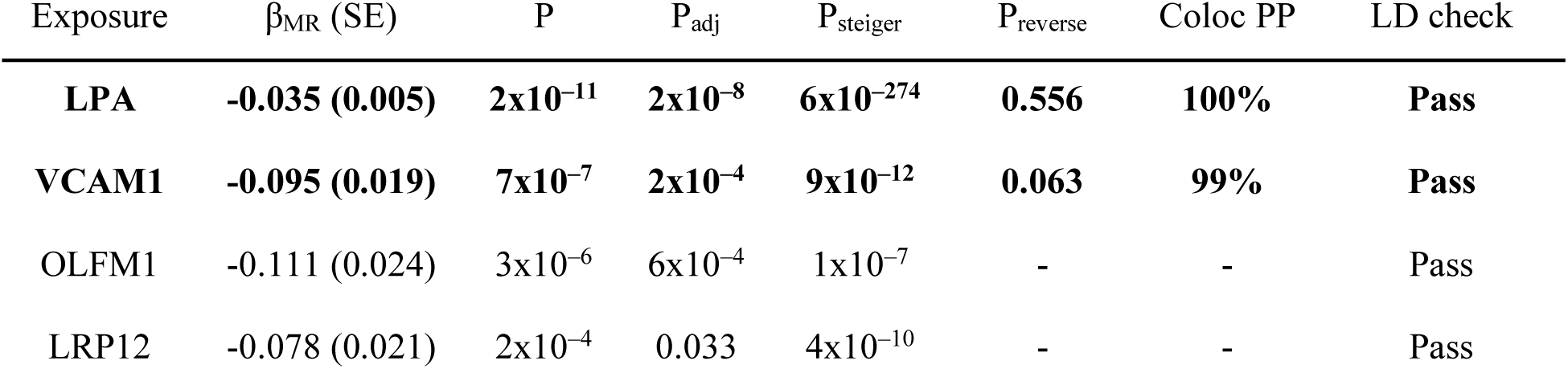
Mendelian randomisation of genetically predicted blood levels of four proteins suggests they have a causal detrimental effect on GIP1. In bold are exposures passing all quality checks, including reverse MR and coloc which required access to full summary statistics. β_MR_—MR effect with standard error in parentheses. P— Nominal P value for the MR effect. P_adj_—FDR-adjusted P value taking into account the 857 proteins tested. P_steiger_—P value for the MR Steiger test assessing if the exposure-outcome pair has the correct causal direction. P_reverse_—P value for the MR effect of GIP1 on the exposure, i.e. evidence of reverse causality. Coloc PP—Posterior probability of colocalisation estimated by coloc. LD check—A secondary check for colocalisation, which requires at least one of the 30 strongest GIP1 SNPs within 500 kb of the pQTL to be in strong LD (r^2^_EUR_ ≥ 0.8) with the pQTL itself.

| Exposure | $\beta_{MR}$ (SE) | P | $P_{adj}$ | $P_{steiger}$ | $P_{reverse}$ | Coloc PP | LD check |
| --- | --- | --- | --- | --- | --- | --- | --- |
| **LPA** | **-0.035 (0.005)** | **$2 \times 10^{-11}$** | **$2 \times 10^{-8}$** | **$6 \times 10^{-274}$** | **0.556** | **100%** | **Pass** |
| **VCAM1** | **-0.095 (0.019)** | **$7 \times 10^{-7}$** | **$2 \times 10^{-4}$** | **$9 \times 10^{-12}$** | **0.063** | **99%** | **Pass** |
| OLFM1 | -0.111 (0.024) | $3 \times 10^{-6}$ | $6 \times 10^{-4}$ | $1 \times 10^{-7}$ | - | - | Pass |
| LRP12 | -0.078 (0.021) | $2 \times 10^{-4}$ | 0.033 | $4 \times 10^{-10}$ | - | - | Pass |

Among the discovered proteins, LPA shows the most significant effect on GIP1 (P_MR_ = 2×10^−8^), with an increase of one standard deviation in genetically predicted blood protein levels causing a decrease of 0.035 (95% CI 0.025–0.045) standard deviations in GIP1. This significance appears to be driven by a consistent detrimental effect across all six GIP1 component traits (β_MR_ range 0.013 to 0.035; all nominal P < 0.05). For a sense of scale, when performing the same MR analysis on the unstandardised parental lifespan GWAS, this equates to a loss of approximately 7 months of life (95% CI 5 months to 9 months) per standard deviation increase in LPA blood levels (**Supplementary Table 6**). The effects of the remaining proteins on GIP1 are larger in magnitude but appear unequally distributed across GIP1 component traits. We estimate a decrease of one standard deviation in genetically predicted VCAM1 blood levels is associated with an increase of approximately 18 months of life (13 months to 23 months); however, comparing VCAM1 effects on standardised GIP1 components, we find its effect to be largest on mid-to-late life ageing traits (lifespan, longevity, resilience) compared to early-life ageing traits (healthspan). Similarly, the effects of OLFM1 and LRP12 appear mostly mediated through late-life ageing traits (resilience and/or longevity). (**Supplementary Figure 2**).

## Discussion

European-ancestry GWAS of healthspan, father and mother lifespan, longevity, frailty, and self-rated health showed genetic correlations between 50% and 96%. Trait-specific genetic correlations indicated some of the differences in genetic architecture may arise from study-specific behavioural associations, such as physical activity and appearance, skin cancers, and daytime napping. As our interest lies in biological ageing, we combined these ageing-related GWAS in a way which maximised power to detect associations with the shared genetic component and downweighed trait-specific genetic associations. The resulting GIP1 trait captured the genetics underlying physical and mental wellbeing, and showed strong inverse genetic correlations with cardiovascular, inflammatory, and neuropsychiatric disease traits. We highlight 27 loci with genome-wide significant effects on GIP1, including two novel loci near *HTT* and *MAML3* that showed directionally consistent evidence of an effect on survival in two additional, independent samples. Across the genome, we found 1,426 genes to be enriched for association with GIP1, which highlighted (among others) the importance of haem metabolism and pathways related to neurogenesis, homeostasis, proteolysis, immunity, and the muscle system in human ageing. Lastly, we performed MR of predicted blood protein levels on GIP1, which revealed the levels of LPA, VCAM1, OLFM1, and LRP12 may be detrimental to multiple indices of healthy ageing.

LPA—apolipoprotein(a)—is a glycoprotein making up the main component of large lipoprotein(a) particles. It is a well-known risk factor for atherosclerotic disease^23^ and is currently being targeted in clinical trials aiming to investigate the effect of LPA-lowering therapy on cardiovascular outcomes^24^. Characterisation of LPA in animal models is limited as the protein is entirely absent from most laboratory animals^23^; however, its high homology to human plasminogen and plasmin suggests it may inhibit fibrinolytic activity, increasing risk of thrombosis^25^. Additionally, human endothelial cell culture experiments show addition of LPA increases cell surface expression of VCAM1^26^ and can increase endothelial cell contraction and permeability^27^. This endothelial dysfunction may mediate its detrimental effect on cardio- and cerebrovascular health.

VCAM1—vascular cell adhesion molecule 1—is a cell adhesion glycoprotein localised predominantly on endothelial cell surfaces. VCAM1 expression is upregulated in response to inflammatory signals and mediates adhesion and transduction of leukocytes across endothelial walls^28^. The link we established between VCAM1 and human ageing relied on a trans-pQTL instrument shared with B2M and is therefore more susceptible to horizontal pleiotropy. That is, the genetic variant may influence VCAM1 levels indirectly, and its effect on human ageing traits could be caused by factors independent of VCAM1 levels. However, only VCAM1 colocalised with the pQTL signal, and experimental evidence from mouse studies suggests the effect of VCAM1 on ageing is likely to be causal. Specifically, VCAM1 levels in blood are known to increase with age in both humans and mice^29^ and treatment with anti-VCAM1 antibodies or an inducible deletion of *Vcam1* improves cognitive performance of aged mice^29^. Of note, similar results have been found for B2M abundance and mouse knockouts^30^, and as such, identification of robust genetic instruments for B2M levels is also warranted.

We were unable to robustly assess colocalisation and reverse causality for the LRP12 (LDL Receptor Related Protein 12) and OLFM1 (Olfactomedin-1) signals as association summary statistics were not available, so the effects of these proteins on human ageing should be interpreted with additional caution. LRP12 belongs to the LDL receptor superfamily and may play a role in brain development^31^ and both tumour proliferation and suppression, depending on the tissue^32,33^. However, the details of its molecular function and its role in the blood remain poorly characterised. Similarly, OLFM1 is a glycoprotein involved in neuronal development and maintenance^34^ which has also been shown to suppress colorectal tumour metastasis^35^. In mouse models, *Olfm1*-knockouts showed reduced cerebral infarction and fertility^36^, but it is unclear if reduction of OLFM1 blood levels will have the same effect in humans.

Importantly, the Mendelian randomisation analysis was restricted to blood pQTL only, which limited our ability to detect causal effects of proteins levels on healthy ageing in other tissues. While it is likely there are proteins with tissue-specific effects on ageing, particularly in the brain, the samples needed for detection of such pQTL are less readily available and therefore more difficult to study at scale (although progress is being made^37^). However, the blood may be a particularly suitable tissue to identify ageing-related proteins. Connecting the circulatory systems of two mice of different ages has shown that systemic factors in the blood can accelerate signs of ageing in the brain, muscle, and liver of the young mouse and can reverse similar signs in the old mouse^38,39^. Likewise, the detrimental cognitive effect of injection of old blood in mice is counteracted when anti-VCAM1 antibodies are concomitantly injected^29^. As such, the blood currently remains one of the most promising tissues to detect ageing-related proteins.

Haem metabolism and iron levels were previously highlighted to have a putative role in human ageing^11^, and here we identify the same pathway using new methods and additional data. Interestingly, both the haem pathway and the proteins we uncovered using MR are strongly linked to vascular and endothelial damage^40^. Across the genome we also found an enrichment for brain tissues and pathways related to neuronal integrity. Given endothelial cells are central to both the cardiovascular system and the blood-brain barrier^41,42^, and endothelial function declines with age^43^, progressive endothelial dysfunction may manifest itself as an age-related disease. Indeed, recent findings suggest the detrimental effects of the APOE*e4 allele—the largest genetic determinant of human ageing—are mediated by an accelerated breakdown of the blood-brain barrier, independently from amyloid-β and tau accumulation^44^. We therefore speculate that molecules involved in maintaining or repairing endothelial integrity may be key to avoiding both age-related cardiovascular injury and neurodegeneration, and recommend further research into this area.

Our study demonstrates that GIP analysis of genetically correlated GWAS can increase power to detect shared genetic architecture and can maximise the heritability of the combined trait. A secondary advantage of the method is the high stability of the resultant GIP1 GWAS, which appears largely robust to the selection of component traits. However, some sociodemographic elements—inherited from one generation to the next, but not necessarily through genetics— appear to be linked to most core ageing traits, and can therefore not be fully separated out in the GIP analysis. For example, the high genetic correlations between GIP1 and employment type, financial satisfaction, and socioeconomic deprivation may inflate GIP1 heritability when these traits are considered to be completely genetic^10^. Similarly, all ageing-related trait GWAS were performed on European-ancestry individuals—with the largest sample contributions from UK Biobank—which may lead to an overrepresentation of associations with European population-specific or even UK-specific confounders^45^, if shared between the studies. Inclusion of a larger and more ancestrally diverse set of ageing-related GWAS in the construction of GIP1 should further converge this phenotype onto a universal ageing phenotype.

Despite these limitations, modelling the shared genetic component of human ageing proxies has allowed us to downweigh non-biological features and propose pathways and proteins which may causally influence the human ageing process. We share the full GIP1 summary statistics without restrictions^46^ to encourage further MR analysis using other biomarkers, and accelerate the discovery of drug targets able to prolong mental and physical wellbeing throughout life.

## Methods

### Data sources

We searched Pubmed and Google Scholar in April 2020 for genome-wide association studies of ageing measures. We included only studies for which we could obtain full summary statistics measured in at least 10,000 European-ancestry individuals. If multiple studies were performed on similar traits we kept the study with the largest sample size. Studies meeting inclusion criteria included GWAS of extreme longevity^16^ (survival past 90th percentile), father and mother lifespan^4^, healthspan^2^, self-reported health^15^, frailty index^8^, epigenetic age acceleration^18^, telomere length^5^, mosaic loss of Y chromosome^7^, and perceived age^17^. As the self-reported health GWAS was performed on the first release of UK Biobank data (N ∼ 150,000), we looked up the same phenotype in the Neale Lab GWAS collection (N ∼ 500,000), which had a larger sample size but was otherwise measured identically^47^. The original derivation of each set of summary statistics is briefly described in the **Supplementary Note**.

For each set of summary statistics, we discarded SNPs that were poorly imputed (INFO < 40%), rare (MAF < 0.5%), or poorly measured (N individuals < 1% of total). The remaining SNPs were aligned to genome build GRCh37 and were harmonised to match UK Biobank SNP IDs (discarding any duplicates). We then estimated the phenotypic variance of the trait (residuals) from independent SNPs using Equation 1 from Winkler *et al*.^48^:

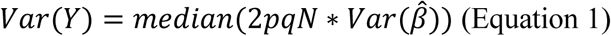

Where *Var*(*Y*) is the phenotypic variance, p and q are the major and minor allele frequencies, N is the total sample size (cases + controls, if applicable), and 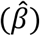 is the variance of the effect size estimate. If N differed by SNP, we calculated the phenotypic variance separately for quintiles of N and took the mean estimate. SNP statistics were then standardised by dividing effect sizes and standard errors by 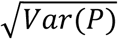.

### Estimation of genetic and non-genetic correlations

The High-definition likelihood (HDL) R package^49^ v1.3.4 was used to calculate SNP heritabilities of ageing-related GWAS and their genetic correlations with each other, using the default European-ancestry LD reference panel and non-MHC SNPs with MAF ≥ 0.01. All GWAS had at least 99.9% of the SNPs in this reference panel, except for the Hannum epigenetic age acceleration GWAS which had 96.29%. Correlations between GWAS due to phenotypic similarity and sample overlap were calculated using Pearson correlation between Z scores of independent SNPs provided by the MultiABEL R package^12^ that were non-significant in both studies (|Z| < 1.96). The number of SNPs used for this calculation is reported in **Supplementary Data 1**.

LD-score regression (LDSC)^50^ v1.0.0 was used to calculate genetic correlations between selected GWAS summary statistics from the GWAS-MAP platform^19^ and the ageing-related GWAS and GIP summary statistics used in our study. The GWAS-MAP platform contains summary statistics for 1,329,912 complex traits and gene expression levels and 3,642 binary traits, derived from UK Biobank^47,51^ and large European-ancestry consortia (e.g. MAGIC, CARDIoGRAM, SSGAC, GIANT) (for an up-to-date list of phenotypes, see https://phelige.com/)^52^. We included all traits which had association statistics for at least 1 million SNPs, measured in at least 10,000 individuals (if continuous) or 2,000 cases and controls (if binary). We further excluded the healthspan and self-rated health GWAS from the GWAS-MAP platform to avoid duplication, after which 728 traits remained. Out of specific interest^53^, we also calculated the genetic correlation between GIPs and a case-control GWAS of COVID-19 hospitalisation, with cases defined as laboratory-confirmed COVID-19 patients experiencing a severe outcome and controls defined as the rest of the population (A2_ALL excluding 23andMe; Release 4)^54^. GWAS with a counterintuitive direction of effect were reversed (e.g. satisfaction traits are coded from high to low in UK Biobank), and each set of GWAS summary statistics were filtered prior to analysis by excluding SNPs located in the Major Histocompatibility Complex, SNPs with low allele frequencies (MAF < 1%) and, if measured, SNPs with suboptimal imputation (INFO < 90%). A full list of 728 GWAS and references can be found in **Supplementary Data 3**. P values were adjusted for multiple testing using Bonferroni correction (729 traits and 6 GIPs).

The same software was used to calculate pairwise correlations between GWAS-MAP statistics, which allowed traits to be clustered based on the magnitude of their genetic similarity. For computational tractability, we only included GWAS that showed large and significant effects on GIP1 (|r_g_| > 0.25; P < 0.05/729/6). Clustering of traits was done hierarchically by maximising the BIC information criterion using mclust v5.4.1, up to a maximum of 100 clusters.

### Identification of shared and unique genetic correlations

For each selected GWAS-MAP phenotype, genetic correlations with ageing-related traits were meta-analysed using a fixed-effect inverse-variance framework, with heterogeneity quantified by the Cochran’s Q and I^2^ statistics as implemented in the meta R package^55^ v4.15-1. GWAS-MAP phenotypes which were significantly correlated with all six ageing-related GWAS at FDR 5% and which did not show substantial evidence of heterogeneity (P_het_ > 0.05 and I^2^ < 50%) were considered to be shared. For GWAS-MAP phenotypes with evidence of heterogeneity, we performed a leave-one-out sensitivity analysis to assess which ageing-related trait(s) contributed most to this heterogeneity. If heterogeneity could be completely removed by excluding a single GWAS (I^2^ = 0%), and exclusion of any other GWAS did not substantially reduce heterogeneity (I^2^ ≥ 50%), the ageing-related trait outlier was considered to have a unique genetic correlation with the GWAS-MAP phenotype.

### Genetically independent phenotype (GIP) analysis of ageing-related trait GWAS

Principal component loadings for six GIP were estimated from the genetic covariance matrix between the six core ageing GWAS, analogous to a principal component analysis of phenotypic correlations. Specifically, eigenvectors from the genetic covariance matrix were transformed into loadings by dividing them by the square root of the phenotypic variance of the GIP. This phenotypic GIP variance was calculated as follows:

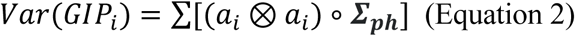

Where *a*_*i*_ is the eigenvector of the *i*-th ageing GIP, and **Σ**_ph_ is the phenotypic variance-covariance matrix of the core ageing traits. As GWAS were standardised, **Σ**_ph_ is equivalent to the phenotypic correlation matrix, with off-diagonal correlations estimated from the correlation observed between independent null Z statistics (described above).

The six core ageing GWAS statistics were then combined on a SNP-by-SNP basis using the principal component loadings to construct genome-wide summary statistics for the six ageing GIPs. GIP effect estimates were calculated by summing effect estimates from the individual ageing-related trait GWAS, each multiplied by their corresponding principal component loading. Standard errors of the GIP effect estimate were calculated by performing the equivalent calculation using variance arithmetic, also taking into account the phenotypic covariance between GWAS:

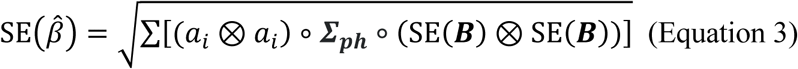

Where SE(***B***) is the vector of SNP standard errors of the core ageing trait effect estimates. Effective sample sizes were then estimated based on the median Z statistic and allele frequencies of null SNPs (|Z| < 1.96), i.e. solving Equation 1 for N. Further technical details of the GIP method are described in the **Supplementary Note**.

GIP summary statistics were calculated for the 7,324,133 SNPs shared between QC’d GWAS statistics, of which 5,353,660 were common (MAF ≥ 5%) and 1,970,474 were rare (MAF < 5%). Finally, standard errors of each GIP GWAS were adjusted to account for the LD-score regression intercept, which ranged from 0.99 (GIP1) to 1.03 (GIP4).

### Association of loci with Finnish and Japanese subject survival

Genome-wide significant loci were defined as 500 kb regions centred on a lead genome-wide significant SNP (P < 5×10^−8^/6) in linkage equilibrium (r^2^ < 0.1) with other lead locus SNPs. Loci were considered to be previously replicated if they had been associated at genome-wide significance with one of the core ageing traits, and also had evidence of an effect in an independent cohort on the same trait (P < 0.05). We attempted to find additional evidence for an effect on ageing for the GIP1 loci which had not been previously replicated.

Effects were first looked up in a GWAS of survival of FinnGen study participants^56^. This study associated SNPs across the genome with the survival of 218,396 Finnish-ancestry individuals (203,244 censored, 15,152 deceased) using Genetic Analysis of Time-to-Event phenotypes (GATE) v0.40. SNP effects were log hazard ratios, calculated from a mixed effect frailty model which adjusted for sex, genotyping batch, birth year, and the first ten genomic PCs as fixed effects, and cryptic relatedness using the genetic relatedness matrix as random effects.

Analogously, the same SNPs (if polymorphic) were regressed against the survival of 135,983 unrelated, Japanese-ancestry individuals (97,365 censored, 30,976 deceased) from Biobank Japan. Here, a fixed effect Cox proportional hazards model was fitted using the survival R package v2.41:

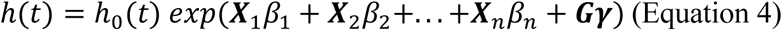

Where h(t) is the hazard at time t, given the subject is alive at time t; h_0_(t) is the baseline hazard at time t; X_1_, X_2_, …, X_n_ are the vectors of covariates with fixed effects β_1_, β_2_, …, β_n_; and ? is the effect of the vector of SNP dosages **G**. All covariates are assumed to cause a proportional change in the hazard, independent of time t. Covariates fitted were sex, disease status, and the first 20 PCs, where disease status refers to one of 47 common diseases in Japan used to recruit the individuals. Each SNP was tested in a separate model.

The SNP effects from both studies were converted from log hazard ratios to approximate years of life by inverting the sign and multiplying the effect estimate and standard errors by ten^3^. For each SNP, a combined effect was calculated by meta-analysing the cohort-specific effects in a fixed-effect framework (weighted using inverse variance), implemented in meta v4.15-1. One-sided P values were adjusted for multiple testing of 23 loci using Bonferroni correction. The collective effect of the 21 loci was calculated using a random-effect framework, to allow for heterogeneity in effect size estimates.

### GIP1 leave-one-out sensitivity analyses

Leave-one-out sensitivity analyses were performed for GIP1, where one at a time, a core ageing trait was excluded and GIP1 loadings and summary statistics were recalculated using the remaining five traits. Genetic and non-genetic correlations were calculated between the original GIP1 and each leave-one-out GIP using HDL inference and null SNP Z statistics, as described above.

To test for heterogeneity in genome-wide significant GIP1 loci, we estimated the difference between the lead SNP effect in GIP1 and the effect in the leave-one-out GIP1 GWAS, taking into account the null correlations between the traits. The standard error of the difference in effects was calculated as follows:

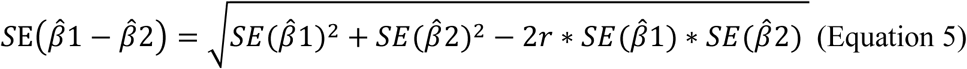

Where is SE(β1) and SE(β2) are the standard errors of the SNP for GIP1 and the leave-one-out GIP1, respectively, and r is the phenotypic correlation between the GWAS. We then used a Wald test to determine the significance of the difference, adjusting P values for multiple testing of 27 loci using Bonferroni correction.

### Lookup of known SNP associations

Lead SNP and close proxies (r^2^_EUR_ ≥ 0.8) of the GIP1 loci were looked up in PhenoScanner^20^ and the GWAS catalog^21^ (accessed 3 December 2020), keeping only the traits with genome-wide significance (P < 5×10^−8^). Triallelic SNPs and any associations with treatments or medications were discarded, before converting associations with the lack of a phenotype into the phenotype itself by inverting the sign (e.g. “Qualifications: none” to “Qualifications”). We then further grouped the traits based on similarities in trait names, keeping the strongest association in the group. This grouping was done by partial matching of trait names—verified manually—and keeping the shortest name. For example, “Melanoma”, “Malignant melanoma”, and “Malignant melanoma of skin” were grouped and renamed to “Melanoma”.

### Tissue enrichment

Stratified LD-score regression v1.0.0 was used to stratify GIP1 SNPs into categories and test whether the proportion of SNP heritability in a category exceeded that expected from the proportion of SNPs in the category^50^. We kept only HapMap3 SNPs, excluding the MHC region and SNPs with MAF < 0.05, and used the 1000 Genomes Phase 3 LD score reference as weights. Categories tested included the 10 groups summarising 220 cell-type specific annotations from Finucane *et al*.^50^, adjusting for the baseline model (v1.2).

### Gene and pathway enrichment

PASCAL^22^ was used to aggregate GIP1 SNP-level P values into gene scores and test these scores for enrichment against predefined gene sets. Gene sets were Hallmark (C1) and Gene Ontology Biological Process (C5.BP) sets from version 7.2 of the Molecular Signatures Database^57^.

GIP1 summary statistics were first aligned to the 1000 Genomes SNP build (matching the PASCAL LD reference), before being tested with default PASCAL parameters, which includes discarding SNPs with MAF < 5% and SNPs in the MHC region. Gene results passing a 5% FDR threshold were considered significant. For each pathway in the C1 and C5.BP datasets, PASCAL calculated two measures of significance based on chi-squared and permutation statistics. We separately adjusted C1 and C5.BP for multiple testing and considered a pathway with both chi-squared and permutation statistics passing a 5% FDR threshold to be significant.

The significant C5.BP pathways were clustered based on their Jaccard similarity coefficient (i.e. size of the intersection of genes divided by the size of the union of genes) using the mclust R package^58^ v5.4.1, which minimised the Bayesian Information Criterion to identify the optimal number of clusters up to a maximum of 100 clusters.

### Mendelian randomisation (MR) of blood protein levels

Genetic instruments for blood protein levels (pQTL) were retrieved from Zheng *et al*.^59^. We included all Tier 1 instruments: cis- and trans-pQTL shown to influence five or fewer proteins (specificity) with no evidence of heterogeneity in effect sizes between multiple protein expression studies (consistency). A total of 857 proteins had non-palindromic SNP instruments (898 total pQTL) present in the GIP1 summary statistics. Two-sample MR of blood protein levels as exposures and GIP1 as outcome was performed using the TwoSampleMR R package^60^ v0.5.5. If multiple pQTL instruments were available for a protein, heterogeneity and MR-Egger sensitivity tests were also performed. This analysis was repeated with the six standardised GIP1 component traits as outcome, as well as the unstandardised, combined parental lifespan GWAS from Timmers *et al*.^4^ to provide an intuitive measure of the effect. The MR effects and standard errors from the latter were multiplied by 10 to convert them from units of negative log hazard ratio to approximate years of life^3^.

GIP1 MR results passing a 5% FDR threshold and sensitivity tests (P_Het_ > 0.05 and P_Egger_ > 0.05; if applicable) were taken forward for follow-up colocalisation tests to rule out LD linkage. For proteins for which we had access to full summary statistics (e.g. LPA, B2M, and VCAM1), we used the coloc R package^61^ v4.0-4 to perform colocalisation analysis using the default parameters, and denoted a posterior colocalisation probability of 80% as evidence of a shared signal. For instruments without full summary statistics, we performed an LD check as described in Zheng *et al*.^59^, which involved checking whether any of the 30 strongest GIP1 SNPs in a 1 Mb region centered on each pQTL were in high LD (r^2^_EUR_ ≥ 0.8) with that pQTL.

Finally, proteins which passed sensitivity and colocalisation tests were subjected to a reverse-causality test using MR-Steiger^62^, and, if full summary statistics were available, a bi-directional MR analysis, both implemented in TwoSampleMR. For the bi-directional MR, we used up to 27 genome-wide significant lead SNPs from GIP1 (shared between GWAS and replacing missing or palindromic SNPs with the next most significant SNP) as instruments and the protein expression statistics as outcome. Proteins which were significant for the MR-Steiger test (P < 0.05) and which did not show evidence of reverse causality in the bi-directional MR (P > 0.05) (if applicable) were considered to show robust evidence of having a causal effect on GIP1.

## Supporting information

Description of supplementary data files

## Data availability

Download links for the 11 ageing-related trait GWAS summary statistics can be found in **Supplementary Table 1**. Summary statistics for GIP1 calculated in this study have been deposited in the Edinburgh DataShare repository, available at https://doi.org/10.7488/ds/2972. Summary statistics of the GWAS-MAP phenotypes used to calculate phenome-wide genetic correlations are available from GeneAtlas (http://geneatlas.roslin.ed.ac.uk/), NealeLab (http://www.nealelab.is/uk-biobank/), or their respective publications. The COVID-19 GWAS summary statistics have been made available by the COVID-19 Host Genetics Initiative at https://www.covid19hg.org/results/. GWAS catalog and PhenoScanner associations can be found at https://www.ebi.ac.uk/gwas/ and http://www.phenoscanner.medschl.cam.ac.uk/, respectively. Curated gene sets (hallmark and gene ontology) are available from the Molecular Signatures Database (https://www.gsea-msigdb.org/). Source data for figures in this study are available in the supplementary documents and upon request from the corresponding author.

## Code availability

HDL: https://github.com/zhenin/HDL/

LDSC: https://github.com/bulik/ldsc/

PASCAL: https://www2.unil.ch/cbg/index.php?title=Pascal

TwoSampleMR: https://mrcieu.github.io/TwoSampleMR/

## Acknowledgements

We would like to acknowledge funding from the Medical Research Council Human Genetics Unit (P.R.H.J.T., J.F.W.: MC_UU_00007/10), the University of Edinburgh (P.K.J.), the Russian Ministry of Education and Science 5-100 Excellence Programme (E.S.T., Y.A.T.) and the Ministry of Education and Science of the RF via the Institute of Cytology and Genetics SB RAS (E.S.T., Y.A.T.: 0259-2021-0009/AAAA-A17-117092070032-4). The FinnGen project and authors (T.T.J.K., W.Z. and A.G.) acknowledge funding from Business Finland (HUS 4685/31/2016 and UH 4386/31/2016) as well as the industry partners AbbVie Inc., AstraZeneca UK Ltd, Biogen MA Inc., Celgene Corporation, Celgene International II Sàrl, Genentech Inc., Merck Sharp & Dohme Corp, Pfizer Inc., GlaxoSmithKline Intellectual Property Development Ltd., Sanofi US Services Inc., Maze Therapeutics Inc., Janssen Biotech Inc, and Novartis AG. The following biobanks are acknowledged for their samples in the FinnGen project: Auria Biobank (www.auria.fi/biopankki), THL Biobank (www.thl.fi/biobank), Helsinki Biobank (www.helsinginbiopankki.fi), Biobank Borealis of Northern Finland (https://www.ppshp.fi/Tutkimus-ja-opetus/Biopankki/Pages/Biobank-Borealis-briefly-in-English.aspx), Finnish Clinical Biobank Tampere (www.tays.fi/en-US/Research_and_development/Finnish_Clinical_Biobank_Tampere), Biobank of Eastern Finland (www.ita-suomenbiopankki.fi/en), Central Finland Biobank (www.ksshp.fi/fi-FI/Potilaalle/Biopankki), Finnish Red Cross Blood Service Biobank (www.veripalvelu.fi/verenluovutus/biopankkitoiminta) and Terveystalo Biobank (www.terveystalo.com/fi/Yritystietoa/Terveystalo-Biopankki/Biopankki/). All Finnish Biobanks are members of BBMRI.fi infrastructure (www.bbmri.fi) and FinBB (https://finbb.fi/). The Framingham Heart Study is funded by National Institutes of Health contract N01-HC-25195, HHSN268201500001I, and 75N92019D00031 (Boston University). This project was funded in part by the Division of Intramural Research, National Heart, Lung, and Blood Institute (NHLBI), National Institutes of Health (NIH), Bethesda, MD. The views expressed in this manuscript are those of the authors and do not necessarily represent the views of the National Heart, Lung, and Blood Institute; the National Institutes of Health; or the U.S. Department of Health and Human Services. For the purpose of Open Access, the authors have applied a CC-BY public copyright licence to any Author Accepted Manuscript version arising from this submission.

## Author Contributions

P.R.H.J.T.: Conceptualization, Methodology, Software, Validation, Formal analysis, Investigation, Writing—Original draft preparation, Writing—Review & editing, Visualization, Project administration. E.S.T.: Formal analysis, Writing— Review & editing. S.S.: Validation, Formal analysis. M.A.: Validation, Formal Analysis. T.T.J.K.: Validation, Formal analysis. W.Z.: Validation, Formal Analysis. S-J.H. Formal analysis. C.Y.: Formal analysis. J.D.: Formal Analysis, Writing— Review & editing. D.L.: Resources, Writing—Review & Editing, Supervision. A.G.: Resources, Supervision. Y.K.: Resources, Data Curation. Y.O.: Resources, Supervision. P.K.J.: Writing—Review & editing, Supervision. J.F.W.: Resources, Writing—Review & editing, Supervision, Funding acquisition. Y.A.T.: Conceptualization, Methodology, Software, Formal analysis, Writing—Review & editing, Supervision.

## References

1. López-Otín, C., Blasco, M. A., Partridge, L., Serrano, M. & Kroemer, G. The hallmarks of aging. Cell 153, 1194 (2013).

2. Zenin, A. et al. Identification of 12 genetic loci associated with human healthspan. Commun. Biol. 2, 41 (2019).

3. Joshi, P. K. et al. Genome-wide meta-analysis associates HLA-DQA1/DRB1 and LPA and lifestyle factors with human longevity. Nat. Commun. 8, 910 (2017).

4. Timmers, P. R. H. J. et al. Genomics of 1 million parent lifespans implicates novel pathways and common diseases and distinguishes survival chances. Elife 8, (2019).

5. Li, C. et al. Genome-wide Association Analysis in Humans Links Nucleotide Metabolism to Leukocyte Telomere Length. Am. J. Hum. Genet. 106, 389–404 (2020).

6. Forsberg, L. A. et al. Mosaic loss of chromosome y in peripheral blood is associated with shorter survival and higher risk of cancer. Nat. Genet. 46, 624–628 (2014).

7. Wright, D. J. et al. Genetic variants associated with mosaic Y chromosome loss highlight cell cycle genes and overlap with cancer susceptibility. Nat. Genet. 49, 674–679 (2017).

8. Atkins, J. et al. A Genome-Wide Association Study of the Frailty Index Highlights Synaptic Pathways in Aging. A Genome-Wide Assoc. Study Frailty Index Highlights Synaptic Pathways Aging 19007559 (2019) doi:10.1101/19007559.

9. Mitnitski, A. B., Mogilner, A. J. & Rockwood, K. Accumulation of deficits as a proxy measure of aging. ScientificWorldJournal. 1, 323–336 (2001).

10. Ruby, J. G. et al. Estimates of the Heritability of Human Longevity Are Substantially Inflated due to Assortative Mating. Genetics 210, 1109–1124 (2018).

11. Timmers, P. R. H. J., Wilson, J. F., Joshi, P. K. & Deelen, J. Multivariate genomic scan implicates novel loci and haem metabolism in human ageing. Nat. Commun. 11, 1–10 (2020).

12. Shen, X. et al. Multivariate discovery and replication of five novel loci associated with Immunoglobulin G N-glycosylation. Nat. Commun. 8, 447 (2017).

13. Ning, Z. et al. Beyond power: Multivariate discovery, replication, and interpretation of pleiotropic loci using summary association statistics. bioRxiv (2015) doi:10.1101/022269.

14. Tsepilov, Y. A. et al. Analysis of genetically independent phenotypes identifies shared genetic factors associated with chronic musculoskeletal pain conditions. Commun. Biol. 3, 1–13 (2020).

15. Harris, S. E. et al. Molecular genetic contributions to self-rated health. Int. J. Epidemiol 46, 994–1009 (2017).

16. Deelen, J. et al. A meta-analysis of genome-wide association studies identifies multiple longevity genes. Nat. Commun. 10, 3669 (2019).

17. Roberts, V., Main, B., Timpson, N. J. & Haworth, S. Genome-Wide Association Study Identifies Genetic Associations with Perceived Age. J. Invest. Dermatol. 140, 2380– 2385 (2020).

18. Gibson, J. et al. A meta-analysis of genome-wide association studies of epigenetic age acceleration. PLoS Genet. 15, e1008104 (2019).

19. Gorev, D. D. et al. GWAS-MAP: a platform for storage and analysis of the results of thousands of genome-wide association scans. 11-Ая Международная Конференция По Биоинформатике Регуляции И Структуры Геномов И Системной Биологии 43–43 (2018) doi:10.18699/bgrssb-2018-020.

20. Kamat, M. A. et al. PhenoScanner V2: an expanded tool for searching human genotype-phenotype associations. Bioinformatics 35, 4851–4853 (2019).

21. Buniello, A. et al. The NHGRI-EBI GWAS Catalog of published genome-wide association studies, targeted arrays and summary statistics 2019. Nucleic Acids Res. 47, D1005–D1012 (2019).

22. Lamparter, D., Marbach, D., Rueedi, R., Kutalik, Z. & Bergmann, S. Fast and Rigorous Computation of Gene and Pathway Scores from SNP-Based Summary Statistics. PLoS Comput Biol 12, e1004714 (2016).

23. Boffa, M. B., Marcovina, S. M. & Koschinsky, M. L. Lipoprotein(a) as a risk factor for atherosclerosis and thrombosis: Mechanistic insights from animal models. Clinical Biochemistry vol. 37 333–343 (2004).

24. Viney, N. J. et al. Antisense oligonucleotides targeting apolipoprotein(a) in people with raised lipoprotein(a): two randomised, double-blind, placebo-controlled, dose-ranging trials. Lancet 388, 2239–2253 (2016).

25. Nordestgaard, B. G. et al. Lipoprotein(a) as a cardiovascular risk factor: Current status. Eur. Heart J. 31, 2844–2853 (2010).

26. Allen, S. et al. Expression of adhesion molecules by Lp(a): a potential novel mechanism for its atherogenicity. FASEB J. 12, 1765–1776 (1998).

27. Cho, T., Jung, Y. & Koschinsky, M. L. Apolipoprotein(a), through its strong lysine-binding site in KIV 10, mediates increased endothelial cell contraction and permeability via a Rho/Rho kinase/MYPT1-dependent pathway. J. Biol. Chem. 283, 30503–30512 (2008).

28. Kong, D. H., Kim, Y. K., Kim, M. R., Jang, J. H. & Lee, S. Emerging roles of vascular cell adhesion molecule-1 (VCAM-1) in immunological disorders and cancer. Int. J. Mol. Sci. 19, 13–17 (2018).

29. Yousef, H. et al. Aged blood impairs hippocampal neural precursor activity and activates microglia via brain endothelial cell VCAM1. Nat. Med. 25, 988–1000 (2019).

30. Smith, L. K. et al. B2-Microglobulin Is a Systemic Pro-Aging Factor That Impairs Cognitive Function and Neurogenesis. Nat. Med. 21, 932–937 (2015).

31. Grote, A. et al. LRP12 silencing during brain development results in cortical dyslamination and seizure sensitization. Neurobiol. Dis. 86, 170–176 (2016).

32. Bethge, N. et al. A gene panel, including LRP12, is frequently hypermethylated in major types of B-cell lymphoma. PLoS One 9, e104249 (2014).

33. Garnis, C., Coe, B. P., Zhang, L., Rosin, M. P. & Lam, W. L. Overexpression of LRP12, a gene contained within an 8q22 amplicon identified by high-resolution array CGH analysis of oral squamous cell carcinomas. Oncogene 23, 2582–2586 (2004).

34. Nakaya, N., Sultana, A., Lee, H. S. & Tomarev, S. I. Olfactomedin 1 interacts with the Nogo A receptor complex to regulate axon growth. J. Biol. Chem. 287, 37171–37184 (2012).

35. Shi, W. et al. Olfactomedin 1 negatively regulates NF-κB signalling and suppresses the growth and metastasis of colorectal cancer cells. J. Pathol. 240, 352–365 (2016).

36. Cheng, A. et al. Pancortin-2 interacts with WAVE1 and Bcl-xL in a mitochondria-associated protein complex that mediates ischemic neuronal death. J. Neurosci. 27, 1519–1528 (2007).

37. Cruchaga, C. et al. Genomic and multi-tissue proteomic integration for understanding the biology of disease and other complex traits. medRxiv 2020.06.25.20140277 (2020) doi:10.21203/rs.3.rs-70284/v1.

38. Rebo, J. et al. A single heterochronic blood exchange reveals rapid inhibition of multiple tissues by old blood. Nat. Commun. 7, 1–11 (2016).

39. Katsimpardi, L. et al. Vascular and neurogenic rejuvenation of the aging mouse brain by young systemic factors. Science (80-.). 344, 630–634 (2014).

40. Balla, J. et al. Haem, haem oxygenase and ferritin in vascular endothelial cell injury. Nephrol. Dial. Transplant. 18, v8–v12 (2003).

41. Higashi, Y. & Yoshizumi, M. Endothelial function. Nippon rinsho. Japanese J. Clin. Med. 61, 1138–1144 (2003).

42. Engelhardt, B. Development of the blood-brain barrier. Cell Tissue Res. 314, 119–129 (2003).

43. Lakatta, E. G. & Levy, D. Arterial and cardiac aging: Major shareholders in cardiovascular disease enterprises: Part I: Aging arteries: A ‘set up’ for vascular disease. Circulation 107, 139–146 (2003).

44. Montagne, A. et al. APOE4 leads to blood–brain barrier dysfunction predicting cognitive decline. Nature 581, 71–76 (2020).

45. Haworth, S. et al. Apparent latent structure within the UK Biobank sample has implications for epidemiological analysis. Nat. Commun. 10, 333 (2019).

46. Timmers, P. R. H. J. et al. Genetically independent phenotype analysis identifies LPA and VCAM1 as drug targets for human ageing. (2021) doi:https://doi.org/10.7488/ds/2972.

47. Neale, B. UK Biobank — Neale lab. http://www.nealelab.is/uk-biobank/%0Ahttp://www.nealelab.is/uk-biobank (2018).

48. Winkler, T. W. et al. Quality control and conduct of genome-wide association meta-analyses. Nat. Protoc. 9, 1192–1212 (2014).

49. Ning, Z., Pawitan, Y. & Shen, X. High-definition likelihood inference of genetic correlations across human complex traits. Nat. Genet. 52, 859–864 (2020).

50. Finucane, H. K. et al. Partitioning heritability by functional annotation using genome-wide association summary statistics. Nat. Genet. 47, 1228–1235 (2015).

51. Canela-Xandri, O., Rawlik, K. & Tenesa, A. An atlas of genetic associations in UK Biobank. Nat. Genet. 50, 1593–1599 (2018).

52. Shashkova, T. I. et al. PheLiGe: an interactive database of billions of human genotype– phenotype associations. Nucleic Acids Res. 49, D1347–D1350 (2021).

53. Ying, K. et al. Genetic and Phenotypic Evidence for the Causal Relationship Between Aging and COVID-19. medRxiv 2020.08.06.20169854 (2020) doi:10.1101/2020.08.06.20169854.

54. Initiative, T. C.-19 H. G. The COVID-19 Host Genetics Initiative, a global initiative to elucidate the role of host genetic factors in susceptibility and severity of the SARS-CoV-2 virus pandemic. Eur. J. Hum. Genet. 28, 715–718 (2020).

55. Schwarzer, G., Carpenter, J. R. & Rücker, G. Meta-Analysis with R. (Springer International Publishing, 2015). doi:10.1007/978-3-319-21416-0.

56. Dey, R. et al. An efficient and accurate frailty model approach for genome-wide survival association analysis controlling for population structure and relatedness in large-scale biobanks. bioRxiv 2020.10.31.358234 (2020) doi:10.1101/2020.10.31.358234.

57. Liberzon, A. et al. The Molecular Signatures Database Hallmark Gene Set Collection. Cell Syst. 1, 417–425 (2015).

58. Scrucca, L., Fop, M., Murphy, T. B. & Raftery, A. E. Mclust 5: Clustering, classification and density estimation using Gaussian finite mixture models. R J. 8, 289–317 (2016).

59. Zheng, J. et al. Phenome-wide Mendelian randomization mapping the influence of the plasma proteome on complex diseases. Nat. Genet. 52, 1122–1131 (2020).

60. Hemani, G. et al. The MR-base platform supports systematic causal inference across the human phenome. Elife 7, e34408 (2018).

61. Giambartolomei, C. et al. Bayesian Test for Colocalisation between Pairs of Genetic Association Studies Using Summary Statistics. PLoS Genet. 10, e1004383 (2014).

62. Hemani, G., Tilling, K. & Smith, G. D. Orienting the causal relationship between imprecisely measured traits using genetic instruments. bioRxiv 13, e1007081 (2017).

