## Supplementary Information for "Genetically independent phenotype analysis identifies LPA and VCAM1 as drug targets for human ageing"

Timmers et al.

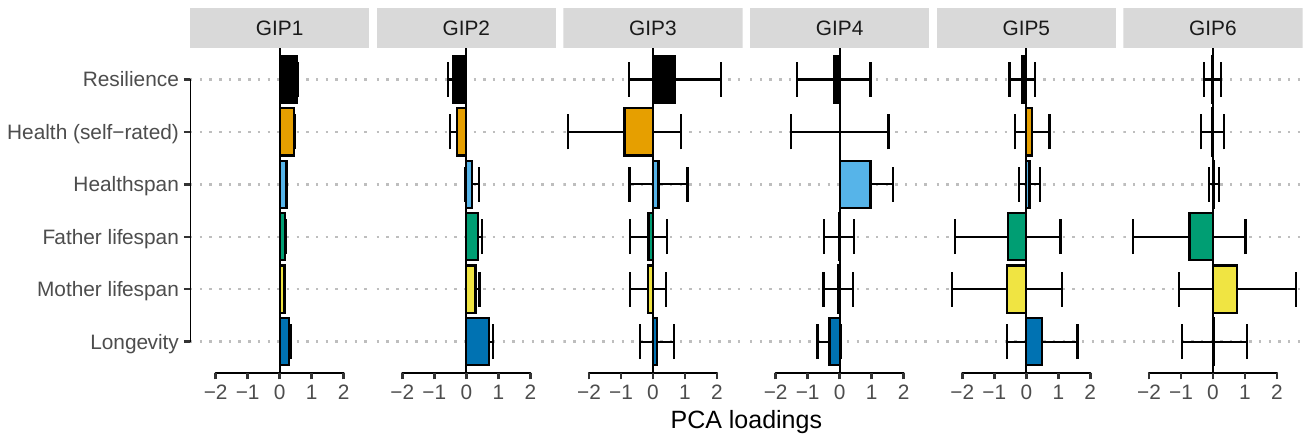

**Supplementary Figure 1: Principal component loadings used to create the genetically independent phenotypes for human ageing.** GIP—Genetically independent phenotype. Resilience—Inverse of frailty index. Error bars represent 95% confidence intervals of the loading estimate.

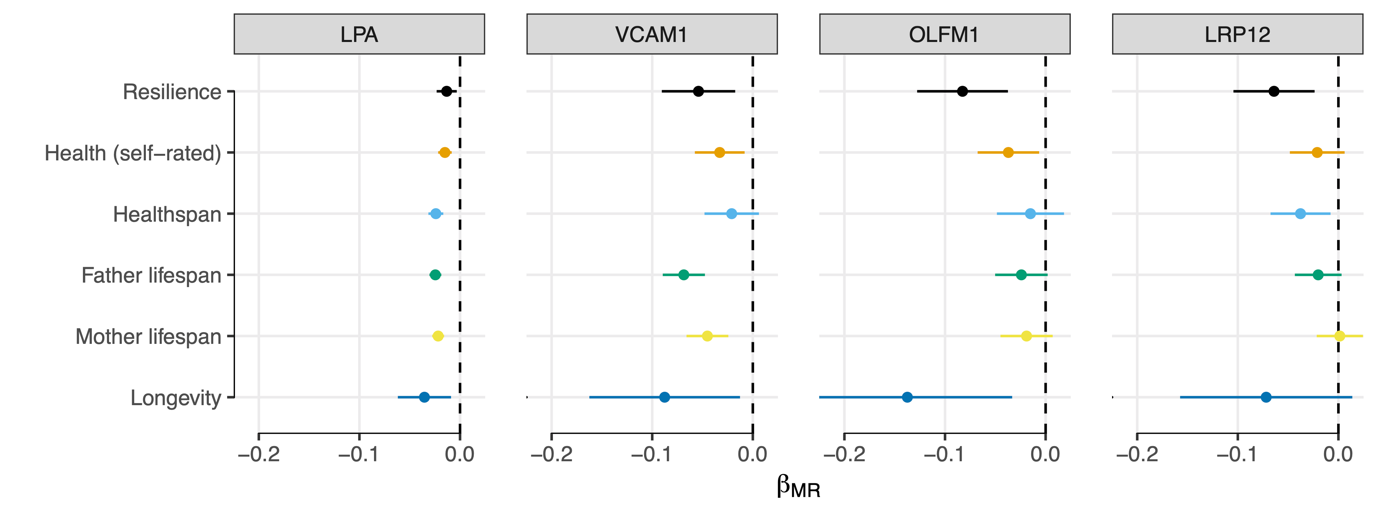

**Supplementary Figure 2: MR of blood protein expression levels on ageing-related GWAS.** Exposures which showed causal evidence of an effect on GIP1 were tested for association with each of the ageing-related traits used to construct GIP1. The x-axis shows the MR effect estimate, with lines representing 95% confidence intervals. LPA—Apolipoprotein(a). VCAM1—Vascular Cell Adhesion Molecule 1. OLFM1—Olfactomedin 1. LRP12— LDL Receptor Related Protein 12.

| Trait | Sample | N | h^2^_SNP_ (SE) | Var(P) | Reference | Source |
| --- | --- | --- | --- | --- | --- | --- |
| Health (self-rated) | UK Biobank | 359,681 | 0.101 (0.002) | 0.53 | Neale Lab 2020 | https://www.dropbox.com/s/ aawh07hlhldbckc/ 2178.gwas.imputed_v3.both_sexes.tsv.bgz |
| Healthspan | UK Biobank | 300,447 | 0.035 (0.002) | 3.56 | Zenin 2019 Comms Bio | https://doi.org/10.5281/ zenodo.1302861 |
| Frailty index | UK Biobank older adults | 164,610 | 0.138 (0.004) | 0.95 | Atkins 2020 medRxiv | https://doi.org/10.6084/ m9.figshare.9204998.v3 |
| Father lifespan | UK Biobank and LifeGen consortium fathers | 492,214 | 0.020 (0.001) | 1.55 | Timmers 2019 eLife | https://doi.org/10.7488/ds/2463 |
| Mother lifespan | UK Biobank and LifeGen consortium mothers | 502,504 | 0.016 (0.001) | 1.92 | Timmers 2019 eLife | https://doi.org/10.7488/ds/2463 |
| Longevity | European Longevity Consortium | 37,082 | 0.071 (0.009) | 7.19 | Deelen 2020 Nat Comms | https://www.longevitygenomics.org/ downloads |
| Perceived age | UK Biobank | 423,992 | 0.073 (0.003) | 4.01 | Roberts 2020 J Invest Derma | https://doi.org/10.5523/ bris.21crwsnj4xwjm2g4qi8chathha |
| EAA (Hannum) | Generation Scotland | 13,493 | 0.144 (0.036) | 28.12 | Gibson 2019 PLOS Gen | https://doi.org/10.1371/ journal.pgen.1008104 |
| EAA (Horvath) | Generation Scotland | 13,493 | 0.101 (0.036) | 20.08 | Gibson 2019 PLOS Gen | https://doi.org/10.1371/ journal.pgen.1008104 |
| Telomere length | ENGAGE consortium and EPIC cohorts | 78,592 | 0.029 (0.007) | 1.04 | Li 2020 Am J Hum Genet | https://doi.org/10.1016/ j.ajhg.2020.02.006 |
| mLOY | UK Biobank men | 85,542 | 0.261 (0.021) | 0.06 | Wright 2017 Nat Gen | https://doi.org/10.1038/ng.3821 |

**Supplementary Table 1: Description of the ten ageing-related trait GWAS used in this study**. N—Number of total individuals used in the study. h^2^_SNP_—SNP-based heritability estimate and standard error (SE), as calculated by High-Definition Likelihood. Var(P)—Phenotypic variance estimated from the GWAS summary statistics. Source—Link to the GWAS article or data repository.

|  |  |  |  |  | Genetic Correlation | |  | Phenotypic Correlation | | |
| --- | --- | --- | --- | --- | --- | --- | --- | --- | --- | --- |
| GIP1 excluding | N_est_ | h^2^_SNP_ | SE |  | Beta | SE |  | Beta | SE | nSNP |
| - | 154,478 | 0.197 | 0.0046 |  | 1.000 | - |  | 1.000 | - | - |
| Resilience | 117,877 | 0.145 | 0.0047 |  | 0.950 | 0.0240 |  | 0.782 | 0.0020 | 52,678 |
| Health (self-rated) | 109,793 | 0.174 | 0.0045 |  | 0.979 | 0.0226 |  | 0.928 | 0.0012 | 53,499 |
| Healthspan | 149,790 | 0.199 | 0.0046 |  | 0.996 | 0.0226 |  | 0.984 | 0.0005 | 53,836 |
| Father lifespan | 154,459 | 0.193 | 0.0046 |  | 0.995 | 0.0220 |  | 0.994 | 0.0003 | 53,997 |
| Mother lifespan | 154,052 | 0.194 | 0.0044 |  | 0.995 | 0.0228 |  | 0.995 | 0.0003 | 54,068 |
| Longevity | 235,354 | 0.162 | 0.0034 |  | 0.987 | 0.0212 |  | 0.725 | 0.0023 | 51,649 |

**Supplementary Table 2: Leave-one-out sensitivity analysis confirms the genetic architecture of GIP1 is stable to the selection of GWAS**. The first row contains data for GIP1 without any of the core ageing trait GWAS excluded. The subsequent rows contain data for GIP1,leaving out the GWAS listed in the first column. N_est_—Number of effective samples, estimated from the median standard error of null SNPs (Z < 1.96). h^2^_SNP_—SNP-based heritability estimate, as calculated by High-Definition Likelihood. SE—Standard error. Beta—Correlation estimate of the leave-one-out GIP1 with the full GIP1. Phenotypic correlation—correlation between the GIP1 and leave-one-out GIP1 Z scores of independent null SNP, capturing both correlations due to phenotypic similarity and sample overlap. nSNP—Number of independent null SNPs used in the phenotypic correlation calculation.

| Nearest gene(s) | rsID | Chr | Position | A1 | A0 | N | Freq1 | Beta1 | SE | P |
| --- | --- | --- | --- | --- | --- | --- | --- | --- | --- | --- |
| NEGR1 | rs2815748 | 1 | 72816147 | G | A | 154,478 | 0.20 | 0.03 | 0.004 | 7E-10 |
| PHTF1 | rs1230682 | 1 | 114293526 | A | G | 154,478 | 0.63 | 0.02 | 0.004 | 4E-10 |
| AFF3 | rs7609078 | 2 | 100490363 | A | G | 154,478 | 0.37 | 0.02 | 0.004 | 3E-10 |
| TRAIP | rs2271961 | 3 | 49878113 | T | C | 154,478 | 0.50 | 0.03 | 0.004 | 8E-14 |
| ADD1 | rs16843603 | 4 | 2928577 | C | T | 154,478 | 0.28 | 0.03 | 0.004 | 1E-10 |
| HTT | rs362273 | 4 | 3227419 | G | A | 154,478 | 0.32 | 0.02 | 0.004 | 4E-10 |
| ANAPC4 | rs34811474 | 4 | 25408838 | A | G | 154,478 | 0.23 | 0.03 | 0.004 | 3E-09 |
| 4q13.2 | rs10434248 | 4 | 67842921 | A | G | 154,478 | 0.57 | 0.02 | 0.004 | 8E-09 |
| MAML3 | rs56172573 | 4 | 140919381 | C | T | 154,478 | 0.38 | 0.02 | 0.004 | 7E-09 |
| C6orf47/GPANK1 | rs805262 | 6 | 31628733 | C | T | 154,478 | 0.52 | 0.03 | 0.004 | 1E-12 |
| HLA-DRB1/DQA1 | rs660895 | 6 | 32577380 | A | G | 154,478 | 0.79 | 0.04 | 0.004 | 1E-22 |
| SLC22A1/A2 | rs9456508 | 6 | 160598596 | T | G | 154,478 | 0.98 | 0.08 | 0.012 | 4E-11 |
| LPA | rs118039278 | 6 | 160985526 | G | A | 154,478 | 0.92 | 0.05 | 0.007 | 1E-11 |
| MAD1L1/SNORA114 | rs11764780 | 7 | 2020904 | C | G | 154,478 | 0.19 | 0.03 | 0.005 | 3E-10 |
| FOXP2 | rs12705966 | 7 | 114248851 | G | A | 154,478 | 0.34 | 0.02 | 0.004 | 7E-10 |
| CSMD3 | rs560719 | 8 | 113032374 | T | C | 154,478 | 0.50 | 0.02 | 0.004 | 5E-09 |
| CDKN2B/-AS1 | rs9632885 | 9 | 22072638 | G | A | 154,478 | 0.52 | 0.02 | 0.004 | 5E-12 |
| CTSF | rs2924807 | 11 | 66341005 | G | C | 154,478 | 0.50 | 0.02 | 0.004 | 5E-09 |
| CCDC90B/DLG2 | rs2512690 | 11 | 83145469 | C | G | 154,478 | 0.34 | 0.02 | 0.004 | 7E-10 |
| TTC12/ANKK1 | rs2186800 | 11 | 113242860 | A | G | 154,478 | 0.54 | 0.02 | 0.004 | 2E-09 |
| USP28/HTR3B | rs61907878 | 11 | 113751052 | C | T | 154,478 | 0.90 | 0.04 | 0.006 | 1E-09 |
| MIR6074 | rs7306710 | 12 | 66376091 | T | C | 154,478 | 0.48 | 0.02 | 0.004 | 4E-10 |
| 12q21.31 | rs6539846 | 12 | 84811217 | A | C | 154,478 | 0.49 | 0.02 | 0.004 | 8E-10 |
| LINC01065 | rs8002970 | 13 | 53924489 | C | G | 154,478 | 0.45 | 0.02 | 0.004 | 6E-11 |
| ZNF652/PHB | rs28394864 | 17 | 47450775 | G | A | 154,478 | 0.54 | 0.02 | 0.004 | 6E-09 |
| APOE | rs429358 | 19 | 45411941 | T | C | 154,478 | 0.84 | 0.07 | 0.005 | 2E-40 |
| ZFP64 | rs67442863 | 20 | 51031131 | T | C | 154,478 | 0.18 | 0.03 | 0.005 | 6E-10 |

| Nearest gene(s) | Resilience | Health (self-rated) | Healthspan | Father lifespan | Mother lifespan | Longevity |
| --- | --- | --- | --- | --- | --- | --- |
| NEGR1 | 0.019 (0.004) | 0.014 (0.003) | 0.011 (0.003) | 0.007 (0.003) | 0.006 (0.002) | 0.021 (0.009) |
| PHTF1 | 0.012 (0.004) | 0.013 (0.002) | 0.007 (0.003) | 0.007 (0.002) | 0.005 (0.002) | 0.023 (0.007) |
| AFF3 | 0.017 (0.004) | 0.012 (0.002) | 0.006 (0.003) | 0.004 (0.002) | 0.004 (0.002) | 0.019 (0.007) |
| TRAIP | 0.019 (0.003) | 0.018 (0.002) | 0.004 (0.003) | 0.002 (0.002) | 0.010 (0.002) | 0.016 (0.007) |
| ADD1 | 0.015 (0.004) | 0.019 (0.003) | 0.006 (0.003) | 0.009 (0.002) | 0.007 (0.002) | 0.015 (0.008) |
| HTT | 0.024 (0.004) | 0.008 (0.003) | 0.007 (0.003) | 0.008 (0.002) | 0.006 (0.002) | 0.011 (0.008) |
| ANAPC4 | 0.020 (0.004) | 0.021 (0.003) | 0.010 (0.003) | 0.002 (0.002) | 0.007 (0.002) | 0.004 (0.011) |
| 4q13.2 | 0.014 (0.004) | 0.011 (0.002) | 0.008 (0.003) | 0.007 (0.002) | 0.005 (0.002) | 0.015 (0.007) |
| MAML3 | 0.011 (0.004) | 0.012 (0.002) | 0.009 (0.003) | 0.007 (0.002) | 0.004 (0.002) | 0.018 (0.007) |
| C6orf47/GPANK1 | 0.023 (0.003) | 0.014 (0.002) | 0.009 (0.003) | 0.005 (0.002) | 0.008 (0.002) | 0.012 (0.009) |
| HLA-DRB1/DQA1 | 0.042 (0.004) | 0.022 (0.003) | 0.016 (0.003) | 0.006 (0.002) | 0.007 (0.002) | 0.014 (0.009) |
| SLC22A1/A2 | 0.045 (0.012) | 0.020 (0.008) | 0.045 (0.009) | 0.031 (0.007) | 0.018 (0.007) | 0.112 (0.029) |
| LPA | 0.015 (0.006) | 0.019 (0.004) | 0.030 (0.005) | 0.031 (0.004) | 0.028 (0.004) | 0.048 (0.017) |
| MAD1L1/SNORA114 | 0.022 (0.004) | 0.020 (0.003) | 0.010 (0.003) | 0.006 (0.003) | 0.005 (0.003) | 0.011 (0.009) |
| FOXP2 | 0.023 (0.004) | 0.011 (0.003) | 0.007 (0.003) | 0.004 (0.002) | 0.005 (0.002) | 0.012 (0.008) |
| CSMD3 | 0.019 (0.003) | 0.007 (0.002) | 0.003 (0.003) | -0.001 (0.002) | -0.002 (0.002) | 0.023 (0.007) |
| CDKN2B/-AS1 | 0.015 (0.003) | 0.005 (0.002) | 0.014 (0.003) | 0.011 (0.002) | 0.007 (0.002) | 0.026 (0.007) |
| CTSF | 0.011 (0.003) | 0.011 (0.002) | 0.003 (0.003) | 0.001 (0.002) | 0.003 (0.002) | 0.027 (0.007) |
| CCDC90B/DLG2 | 0.020 (0.004) | 0.012 (0.002) | 0.007 (0.003) | 0.003 (0.002) | 0.003 (0.002) | 0.015 (0.008) |
| TTC12/ANKK1 | 0.019 (0.003) | 0.015 (0.002) | 0.010 (0.003) | 0.004 (0.002) | 0.005 (0.002) | 0.003 (0.007) |
| USP28/HTR3B | 0.020 (0.006) | 0.018 (0.004) | 0.015 (0.004) | 0.013 (0.003) | 0.014 (0.003) | 0.038 (0.013) |
| MIR6074 | 0.017 (0.003) | 0.013 (0.002) | 0.002 (0.003) | 0.005 (0.002) | 0.007 (0.002) | 0.017 (0.007) |
| 12q21.31 | 0.012 (0.003) | 0.012 (0.002) | 0.008 (0.003) | 0.004 (0.002) | 0.006 (0.002) | 0.021 (0.007) |
| LINC01065 | 0.016 (0.003) | 0.017 (0.002) | 0.004 (0.003) | -0.002 (0.002) | 0.000 (0.002) | 0.021 (0.007) |
| ZNF652/PHB | 0.013 (0.004) | 0.013 (0.002) | 0.010 (0.003) | 0.003 (0.002) | 0.003 (0.002) | 0.015 (0.007) |
| APOE | 0.005 (0.005) | -0.001 (0.003) | 0.007 (0.004) | 0.033 (0.003) | 0.048 (0.003) | 0.190 (0.012) |
| ZFP64 | 0.021 (0.004) | 0.020 (0.003) | 0.007 (0.003) | 0.003 (0.003) | 0.004 (0.003) | 0.018 (0.009) |

**Supplementary Table 3: Genomic loci associated with the first genetic principal component of human ageing (GIP1).** Loci were defined as 500 kb regions centred on a lead genome-wide significant SNP (P < 5x10^–8^/6) in linkage equilibrium (r^2^ < 0.1) with other lead locus SNPs. Nearest gene(s)—Closest genes upstream/downstream to the lead SNP (within 250 kb), or if none, the closest cytogenetic band. rsID—The lead SNP within the locus. Chr—Chromosome. Position—Base-pair position (GRCh37). A1—Effect allele, associated with higher GIP1. A0—Other, non-effect allele. N—Sample size estimated from the GIP1 GWAS standard errors. Freq1—Allele frequency of the effect allele in UK Biobank. Beta1—Effect of the SNP on GIP1 in standard deviation units. SE—Standard error. P—Nominal, Wald-ratio P value. The final columns contain the effects (and standard errors in parentheses) of the SNP on each core ageing traits used to construct GIP1, in standard deviation units.

| Nearest gene(s) | rsID | A1 | N | Dead | Freq1 (FG) | Freq1 (BBJ) | Years1 | SE | P | P_adj_ |
| --- | --- | --- | --- | --- | --- | --- | --- | --- | --- | --- |
| **HTT** | **rs362273** | **G** | **346,737** | **46,128** | **0.44** | **0.32** | **0.211** | **0.0689** | **0.001** | **0.025** |
| **MAML3** | **rs56172573** | **C** | **346,726** | **46,128** | **0.36** | **0.39** | **0.209** | **0.0732** | **0.002** | **0.049** |
| TRAIP | rs2271961 | T | 346,737 | 46,128 | 0.64 | 0.50 | 0.212 | 0.0763 | 0.003 | 0.063 |
| C6orf47/GPANK1 | rs805262 | C | 346,737 | 46,128 | 0.59 | 0.52 | 0.123 | 0.0685 | 0.037 | 0.844 |
| PHTF1 | rs1230682 | A | 346,737 | 46,128 | 0.56 | 0.63 | 0.135 | 0.0767 | 0.039 | 0.898 |
| ADD1 | rs16843603 | C | 346,737 | 46,128 | 0.35 | 0.28 | 0.201 | 0.1167 | 0.042 | 0.971 |
| TTC12/ANKK1 | rs2186800 | A | 346,737 | 46,128 | 0.51 | 0.53 | 0.170 | 0.0989 | 0.043 | 0.995 |
| ZFP64 | rs67442863 | T | 346,737 | 46,128 | 0.16 | 0.19 | 0.150 | 0.0908 | 0.049 | 1.000 |
| NEGR1 | rs2815748 | G | 346,737 | 46,128 | 0.14 | 0.20 | 0.205 | 0.1449 | 0.078 | 1.000 |
| 4q13.2 | rs10434248 | A | 346,726 | 46,128 | 0.53 | 0.56 | 0.092 | 0.0684 | 0.089 | 1.000 |
| ANAPC4 | rs34811474 | A | 218,396 | 15,152 | 0.23 | 0.00 | 0.154 | 0.1421 | 0.140 | 1.000 |
| MAD1L1/SNORA114 | rs11764780 | C | 346,737 | 46,128 | 0.29 | 0.18 | 0.079 | 0.1025 | 0.221 | 1.000 |
| MIR6074 | rs7306710 | T | 346,736 | 46,128 | 0.51 | 0.49 | 0.061 | 0.0862 | 0.239 | 1.000 |
| USP28/HTR3B | rs61907878 | C | 218,396 | 15,152 | 0.90 | 0.00 | 0.087 | 0.2003 | 0.333 | 1.000 |
| CTSF | rs2924807 | G | 346,737 | 46,128 | 0.50 | 0.50 | 0.005 | 0.0683 | 0.473 | 1.000 |
| CSMD3 | rs560719 | T | 346,737 | 46,128 | 0.49 | 0.50 | -0.009 | 0.0678 | 0.554 | 1.000 |
| LINC01065 | rs8002970 | C | 346,737 | 46,128 | 0.45 | 0.45 | -0.034 | 0.0674 | 0.693 | 1.000 |
| CCDC90B/DLG2 | rs2512690 | C | 346,737 | 46,128 | 0.28 | 0.32 | -0.038 | 0.0694 | 0.709 | 1.000 |
| FOXP2 | rs12705966 | G | 346,737 | 46,128 | 0.31 | 0.32 | -0.086 | 0.1218 | 0.761 | 1.000 |
| ZNF652/PHB | rs28394864 | G | 346,737 | 46,128 | 0.51 | 0.54 | -0.070 | 0.0696 | 0.842 | 1.000 |
| SLC22A1/A2 | rs9456508 | T | 346,737 | 46,128 | 0.98 | 0.98 | -0.096 | 0.0903 | 0.857 | 1.000 |
| AFF3 | rs7609078 | A | 346,737 | 46,128 | 0.35 | 0.37 | -0.073 | 0.0678 | 0.859 | 1.000 |
| 12q21.31 | rs6539846 | A | 346,736 | 46,128 | 0.55 | 0.49 | -0.100 | 0.0674 | 0.932 | 1.000 |

**Supplementary Table 4: Association of lead GIP1 SNPs with survival in an independent sample of Finnish and Japanese individuals.** Nearest gene(s)—Closest genes upstream/downstream to the lead SNP (within 250 kb), or if none, the closest cytogenetic band. rsID—The lead SNP within the locus. A1—Effect allele, associated with higher GIP1. Freq1—Allele frequency of the effect allele in FinnGen (FG) or Biobank Japan (BBJ). Years1—Effect of the SNP on survival per A1 allele in approximate years of life. SE—Standard error. P—One-sided P value for a directionally consistent effect with GIP1. P_adj_—Bonferroni-adjusted P value, taking into account 23 tests. Both cohorts tested rs16891156 as a proxy for the lead GIP1 SNP near *SLC22A1/A2*, and Biobank Japan used rs494865 and rs11523976 as proxies for *CTSF* and *CCDC90B/DLG2*, respectively. All of these proxies were in perfect linkage disequilibrium (r^2^ = 1.0) with the lead GIP1 SNP for their respective ancestries.

|  | Proportion | |  | Enrichment | | |  | Model coefficients | | | | |
| --- | --- | --- | --- | --- | --- | --- | --- | --- | --- | --- | --- | --- |
| Tissue | SNPs | h^2^ (SE) |  | Beta | SE | P |  | Beta | SE | Z | P | FDR |
| **CNS** | **0.15** | **0.42 (0.03)** |  | **2.85** | **0.19** | **5E-19** |  | **5E-08** | **7E-09** | **6.19** | **6E-10** | **6E-09** |
| **Adrenal_Pancreas** | **0.09** | **0.31 (0.03)** |  | **3.31** | **0.34** | **2E-10** |  | **4E-08** | **1E-08** | **2.86** | **4E-03** | **4E-02** |
| GI | 0.17 | 0.31 (0.03) |  | 1.84 | 0.19 | 3E-05 |  | -2E-08 | 9E-09 | -2.42 | 2E-02 | 2E-01 |
| Kidney | 0.04 | 0.16 (0.02) |  | 3.72 | 0.50 | 1E-07 |  | 3E-08 | 2E-08 | 1.83 | 7E-02 | 7E-01 |
| Hematopoietic | 0.23 | 0.42 (0.03) |  | 1.81 | 0.15 | 1E-07 |  | 1E-08 | 8E-09 | 1.59 | 1E-01 | 1E+00 |
| SkeletalMuscle | 0.10 | 0.25 (0.03) |  | 2.39 | 0.26 | 2E-07 |  | -1E-08 | 1E-08 | -1.40 | 2E-01 | 1E+00 |
| Other | 0.20 | 0.38 (0.04) |  | 1.88 | 0.19 | 9E-06 |  | -1E-08 | 9E-09 | -1.27 | 2E-01 | 1E+00 |
| Cardiovascular | 0.11 | 0.28 (0.03) |  | 2.52 | 0.29 | 4E-07 |  | -1E-08 | 1E-08 | -1.18 | 2E-01 | 1E+00 |
| Liver | 0.07 | 0.20 (0.02) |  | 2.79 | 0.33 | 1E-07 |  | 1E-08 | 1E-08 | 0.95 | 3E-01 | 1E+00 |
| Connective_Bone | 0.11 | 0.27 (0.03) |  | 2.33 | 0.25 | 2E-07 |  | -5E-09 | 1E-08 | -0.49 | 6E-01 | 1E+00 |

**Supplementary Table 5: Enrichment of GIP1 SNP heritability within the central nervous system and adrenal/pancreas tissues.** Tissues with model coefficients passing a 5% FDR threshold are highlighted in bold. SE—Standard Error. CNS—Central Nervous System.

|  | Standard deviation units | | | | | |  | Years |
| --- | --- | --- | --- | --- | --- | --- | --- | --- |
| Exposure | Resilience | Health (self-rated) | Healthspan | Father lifespan | Mother lifespan | Longevity |  | Parental Lifespan |
| LPA | -0.013 (0.005) | -0.015 (0.003) | -0.024 (0.004) | -0.025 (0.003) | -0.022 (0.003) | -0.035 (0.013) |  | -0.61 (0.06) |
| VCAM1 | -0.054 (0.019) | -0.033 (0.013) | -0.021 (0.014) | -0.069 (0.011) | -0.045 (0.011) | -0.088 (0.038) |  | -1.47 (0.21) |
| OLFM1 | -0.083 (0.023) | -0.037 (0.016) | -0.015 (0.017) | -0.024 (0.013) | -0.019 (0.013) | -0.137 (0.053) |  | -0.59 (0.26) |
| LRP12 | -0.064 (0.021) | -0.021 (0.014) | -0.038 (0.015) | -0.020 (0.012) | 0.001 (0.012) | -0.072 (0.044) |  | -0.32 (0.23) |

**Supplementary Table 6: MR of blood protein expression levels on ageing-related GWAS.** Exposures which showed causal evidence of an effect on GIP1 were tested for association with each of the ageing-related traits used to construct GIP1 (standardised units), as well as the unstandardised GWAS meta-analysis of father and mother lifespan (in units of approximate years of life). Displayed are the MR effects with standard errors in parentheses. LPA—Apolipoprotein(a). VCAM1—Vascular Cell Adhesion Molecule 1. OLFM1—Olfactomedin 1. LRP12— LDL Receptor Related Protein 12.

### Supplementary Note

#### Description of ageing-related trait GWAS summary statistics

We provide a short description of the derivation of each set of ageing-related GWAS summary statistics used in our study. Further methodological detail can be found in the original studies.

##### Longevity

Deelen *et al.*^1^ assessed the genetics of reaching an exceptional age compared to population- and sex-specific mortality rates. Individuals whose age at last contact (or age at death) was above the 90th percentile were classified as cases (N = 11,262), while those below the 60th percentile were classified as controls (N = 25,483). Summary statistics on 1000G Phase 1 version 3 SNPs were calculated separately for 18 European-ancestry cohorts (birth years 1875 to 1990) by fitting a logistic regression model, adjusting for clinical site, cryptic relatedness, and/or the first four genetic principal components. Cohort-specific statistics were then combined in a fixed-effect meta-analysis, weighted by the effective sample size of each cohort.

After applying our quality control filters (INFO ≥ 40%, MAF ≥ 0.5%, measured in at least 1% of total sample, and no duplicate SNP IDs), summary statistics for 6,656,519 common (MAF ≥ 5%) and 2,094,480 rare (MAF < 5%) SNPs remained.

##### Father and mother lifespan

Timmers *et al.*^2^ assessed the genetics of parent survival of unrelated individuals from UK Biobank and 26 other European-ancestry cohorts. The lifespans of 500,193 fathers and 512,047 mothers aged 40 or older were included, 60% of which had died. Summary statistics on HRC-imputed SNPs were calculated separately for father and mother survival in each cohort. Specifically, parental survival was modelled from parent age and dead/alive status using Cox proportional hazards models, adjusting for subject sex, study-specific covariates (e.g. genotyping batch), and at least 10 genetic principal components. Martingale residuals of these survival models were used as phenotype in a linear regression model testing for association with SNP dosage. Cohort-specific statistics were combined in a fixed-effect meta-analysis, separately for father and mother survival.

After applying our quality control filters, father lifespan summary statistics contained 5,553,356 common (MAF ≥ 5%) and 3,360,107 rare (MAF < 5%) SNPs, while mother lifespan summary statistics contained 5,437,832 common and 3,478,109 rare SNPs.

##### Healthspan

Zenin *et al.*^3^ assessed the genetics of healthspan: the number of years until the first reported hospital incidence of one of seven age-related diseases (cancer, diabetes, myocardial infarction, stroke, chronic obstructive pulmonary disease, dementia, and congestive heart failure) or a reported incidence of death. Summary statistics on HRC-imputed SNPs were calculated for the healthspans of 300,477 unrelated, British-ancestry individuals from UK Biobank (birth years ~1934 to ~1971), 28.3% of which were classified as no longer healthy. This was done by modelling healthspan using individuals’ age and disease status in a Cox-Gompertz proportional hazards model, adjusting for sex, assessment centre, genotyping batch, and the first 40 genetic principal components. Martingale residuals from this model were then regressed against SNP dosage.

After applying our quality control filters, summary statistics for 5,429,270 common (MAF ≥ 5%) and 3,326,508 rare (MAF < 5%) SNPs remained.

##### Frailty index

Atkins *et al.*^4^ assessed the genetics of frailty, calculated as an integer score based on the self-reported presence or absence of 49 markers of physical and mental deficiency. These markers included sensory deficits (glaucoma, cataract, hearing difficulties), psychological deficits (fatigue, depression, loneliness, etc), infirmity (falls & fractures), cancers, pain (including migraine), as well as a variety of cardiometabolic, musculoskeletal, immunological, and gastrointestinal disorders and disease markers^5^. Summary statistics on HRC-imputed SNPs were calculated for 164,610 older individuals aged between 60 and 70 years, which had frailty indices between 0 and 27. Specifically, the proportion of deficits was quantile-normalised and then regressed against SNP dosage in a linear model, adjusting for sex, age, assessment centre, genotyping array, and cryptic relatedness.

After applying our quality control filters, summary statistics for 6,914,134 common (MAF ≥ 5%) and 4,129,439 rare (MAF < 5%) SNPs remained.

##### Self-reported health

Harris *et al.*^6^ assessed the genetics of self-reported health in UK Biobank individuals from the first release of genetic information (N = 111,749). After their study was published, a GWAS of the same phenotype (UKBB field 2178) was performed by NealeLab (<http://www.nealelab.is/uk-biobank>) on the full UK Biobank sample (birth years ~1934 to ~1971), including only unrelated European-ancestry individuals (N = 359,681). Given the larger sample size, we chose to use the NealeLab GWAS, but mention the Harris study as it provides a peer-reviewed discussion of the trait and its limitations. Overall health rating was assessed in UK Biobank using a touch screen questionnaire asking ‘In general how would you rate your overall health?’, which quantified responses as a categorical variable with values Excellent, Good, Fair, and Poor. Summary statistics on HRC-imputed SNPs were calculated using PHEASANT^7^, fitting an ordinal logistic regression model adjusted for age, age^2, sex, interactions between age/age^2 and sex, as well as the first 20 genetic principal components.

After applying our quality control filters, summary statistics for 6,949,308 common (MAF ≥ 5%) and 3,883,623 rare (MAF < 5%) SNPs remained. Additionally, we reversed the direction of the genetic effects to provide a more intuitive scale, with positive SNP coefficients indicating better self-rated health.

##### Epigenetic age acceleration (EAA)

Gibson et al.^8^ assessed the genetics of EAA, calculated as the residual when regressing epigenetic age (age predicted from epigenetic markers) against chronological age. Positive EAA thus indicates an individual is predicted to have an epigenetic age which exceeds their chronological age. Two models were used to calculate epigenetic age: the first uses 71 CpG sites differentially methylated with age in whole blood (Hannum), and the second uses 353 CpG sites with weights derived from 51 different tissues, adjusted for blood cell counts (Horvath). The sample included 12 European-ancestry datasets consisting of 13,493 individuals aged 10 to 98 (mean age 57.5), the largest of which was Generation Scotland (N = 5,100; mean Hannum EAA –0.0014 years [SD 3.82]; mean Horvath EAA –0.0014 years [SD 4.24]). Summary statistics on HRC-imputed SNPs (Generation Scotland) or 1000G Phase 1 imputed SNPs (other cohorts) were calculated using linear regression of epigenetic age acceleration against SNP dosage, adjusted for sex. Depending on the cohort, relatedness and population structure were adjusted for by fitting genetic principal components as fixed effects and/or genomic relationship matrices as random effects. Cohort-specific statistics were then combined in a fixed-effect inverse-variance meta-analysis.

After applying our quality control filters, Hannum EAA summary statistics consisted of 5,094,910 common (MAF ≥ 5%) and 836,261 rare (MAF < 5%) SNPs, while Horvath summary statistics consisted of 5,101,490 common and 830,617 rare SNPs.

##### Mosaic loss of Y chromosome (mLOY)

Wright *et al.*^9^ assessed the genetics of cellular loss of the Y chromosome in the whole blood of UK Biobank men of white European ancestry. mLOY values were estimated for 67,034 males using the normalised signal intensity (mean LLR) of SNPs on the male-specific region of the Y chromosome (mean LLR 0.00 [SD 0.067]). Summary statistics on HRC-imputed SNPs were calculated using a mixed model framework, which adjusted for sex and genotyping array using fixed effects and fitted a random effect to a genetic relatedness matrix to account for population stratification and cryptic relatedness.

After applying our quality control filters, summary statistics for 6,958,267 common (MAF ≥ 5%) and 4,159,385 rare (MAF < 5%) SNPs remained.

##### Perceived age

Roberts *et al.*^10^ assessed the genetics of perceived age. The phenotype was measured as a categorial variable in 423,992 individuals from UK Biobank (primarily but not exclusively from European ancestries) using an online questionnaire. Participants reported being perceived by others as older than their age (N = 8,630), about their age (N = 103,300), or younger than their age (N = 312,062), which was quantified as 0, 0.5, and 1, respectively. Summary statistics on HRC-imputed SNPs were calculated by regressing SNP dosage against perceived age values in a linear mixed model framework, with age, sex, and assessment centre as fixed effects and the genetic relationship matrix as a random effect.

After applying our quality control filters, summary statistics for 5,425,902 common (MAF ≥ 5%) and 3,286,322 rare (MAF < 5%) SNPs remained. Additionally, we reversed the direction of the genetic effects to provide a more intuitive scale, with positive SNP coefficients indicating a higher perceived age.

##### Telomere length

Li *et al.*^11^ assessed the genetics of telomere length, measured from leukocyte DNA using quantitative polymerase chain reaction. The phenotype was a Z-transformed ratio of the telomere repeat number divided by the number of single gene repeats. The sample included 78,592 unrelated European-ancestry individuals aged 12 to 99, from 21 cohorts within the ENGAGE consortium (N = 46,898) as well as the EPIC InterAct and EPIC CVD studies (N = 31,694; stratified by genotyping platform and disease status). Summary statistics on HRC-imputed SNPs were calculated by regressing SNP dosage against the phenotype, adjusting for subject sex and age, as well as study-specific covariates (e.g. genotyping batch, top genetic principal components, family structure). Cohort-specific statistics were then combined largely using a fixed-effect inverse-variance meta-analysis framework.

After applying our quality control filters, summary statistics for 6,490,990 common (MAF ≥ 5%) and 4,182,895 rare (MAF < 5%) SNPs remained.

#### Genetically independent phenotype (GIP) analysis methodology

To decompose the traits of interest into genetically independent components, we use the matrix of genetic covariances Ω (instead of the matrix of phenotypic covariance used in conventional principal component analysis (PCA) for biological traits) for extraction of eigenvectors *a_i_* (vectors of orthogonal transformation coefficients). The resulting principal components are referred to as “genetically independent phenotypes” (GIPs).

Technical details of the genetic principal component analysis approach are provided below.

Denote the following variables:

Ω – the matrix of genetic covariances (*m* x *m*, where *m* is the number of traits)

Ω_SE_ – the matrix of the standard errors of genetic covariances (*m* x *m*)

∑_ph_ – the matrix of phenotypic covariances (*m* x *m*); in the case of standardized traits it is equal to the matrix of phenotypic correlations

B – the matrix of effect sizes (*β*) for *m* phenotypes (*M* x *m*, where *M* is the number of SNPs in the analysis). *b_i_* is the *i*-th column of B.

SE – the matrix of standard errors of *β* for *m* phenotypes (*M* x *m*). *SE*_i_ is the *i*-th column of SE.

*varY*_i_ – the variance of the *i*-th trait. After standardization, *varY*_i_ = 1.

*SD_i_* – the standard deviation of *i*-th trait. ${SD}_{i}= \sqrt{{varY}_{i}}$

B_s_ – the matrix of standardized *β* for *m* phenotypes (*M* x *m*). b^s^_i_ is the *i*-th column of B_s_.

SE_s_ – the matrix of standardized standard errors for *m* phenotypes (*M* x *m*). *SE^s^_i_* is the *i*-th column of SE_s_.

A – the matrix of eigenvectors of Ω (*m* x *m*). Each column is *a_i_*, {*a_1_…a_m_*} – the vector of orthogonal transformation coefficients of *m* original traits into *m* GIPs {GIP_1_.. GIP_m_}

A_s_ – the matrix of scaled eigenvectors of Ω (*m* x *m*). Each column is *a^s^_i_*, {*a^s^_1_*…*a^s^_m_*} – the vector of orthogonal transformation coefficients of *m* original traits into *m* GIPs scaled to make the GIPs’ variance equal to 1.

*L* – the vector of eigenvalues {*l_1_…l_m_*}.

We performed the following procedure to calculate GIPs for studied phenotypes:

1. Estimated Ω and Ω_SE_.
2. Estimated *varY*_i_ and Pearson correlation matrix for phenotypes.
3. Estimated standardized effect sizes and standard errors from Z-scores (${SE}_{i}^{s}=\sqrt{1/{({Z_{i}}^{2}+N)}}$ and $\beta_{i}^{s}=Z_{i}*{SE}_{i}^{s}$), where *N* is sample size.
4. Checked whether all eigenvalues were positive for Ω.
5. Estimated eigenvalues (L) and the matrix of eigenvectors (A) of Ω.
6. If the coefficient of a given eigenvector for first phenotype was negative ($a_{i,trait\_1}$< 0), we changed the signs for all coefficients in its eigenvector ($a_{i}=-a_{i}$)
7. Estimated variance for GIP as ${var(GIP}_{i})=\sum\left[ (a_{i}\bigotimes a_{i})\circ\sum_{ph} \right]$, where $\bigotimes$ is an outer product.
8. Scaled coefficients for GIPs as $a_{i}^{s}={a_{i}}/{{SD(GIP}_{i})}$.
9. Estimated 95% CI for *a_i_* (see below)^*^.
10. Provided GWAS results for each GIP (see below)^**^.

^*^For estimation of 95% CI for GIPs, the Monte Carlo approach was used. We performed 1000 cycles of simulations. In each round, we simulated the noise component for matrix of genetic correlations Ω - the matrix Ω*^noise^* (*m* x *m*). Each element *i,j* (*i* > *j*) Ω*^noise^* is sampled from the normal distribution with zero mean and standard deviation equal to the *i,j* element of matrix of standard errors ($\Omega_{i,j}^{SE}$). The resulting covariance matrix was obtained as the sum of Ω and Ω*^noise^*. Than the standardized eigenvalues were calculated as described above formulating the empirical distribution of each element of A_s_ matrix. For each element of A_s_, 95% CI was obtained as an absolute difference between 0.975 and 0.025 quantiles divided by 2.

^**^Estimation of GWAS for GIPs was performed using the following procedure:

1. Unscaled effect sizes for *M* SNPs were calculated as $\beta_{{GIP}_{i},unscaled}=B_{s}\times a_{i}^{s}$, where $\times$ is an inner product.
2. Estimation of variance for GIPs was calculated as ${varGIP}_{i}=\sum\left[ (a_{i}\bigotimes a_{i})\circ\sum_{ph} \right]$. Phenotypic correlation matrix $\sum_{ph}$ was obtained from the specific studied population.
3. The unscaled standard errors of effect sizes for *M* SNPs were calculated as

${SE}_{{GIP}_{i},unscaled}=\sqrt{\sum\left[ (a_{i}\bigotimes a_{i})\circ\sum_{ph}\circ({SE}_{i}^{s}\bigotimes{SE}_{i}^{s}) \right]}$.

4. Scaling of effect sizes and standard errors were performed by dividing of unscaled effect sizes and standard errors by $\sqrt{{varGIP}_{i}*2*p_{i}*(1-p_{i})}$, where $p_{i}$ is allele frequency for *i*th SNP.
5. The corresponding *P*-values were estimated using Wald test ($Z-score={\beta_{{GIP}_{i}}}/{{SE}_{{GIP}_{i}}}$).
6. The estimation of effective sample size for each GIP was performed as $N_{{GIP}_{i}}=median(1/{({{SE}_{{GIP}_{i},unscaled}}^{2})})$

The total genetic variance of m original traits explained by each GIP was calculated as $R_{{GIP}_{i}}^{2}={l_{i}}/{\sum_{i=1}^{m} l_{i}}$.

The heritability of each GIP was calculated as $h_{{GIP}_{i}}^{2}=\frac{\sum\left[ (a_{i}\bigotimes a_{i})\circ\Omega\right]}{\sum\left[ (a_{i}\bigotimes a_{i})\circ\sum_{ph} \right]}$

Genetic correlations between GIPs and original traits as well as between each other were calculated as

$$\rho_{genetic}\left\{ c_{1} | c_{2} \right\}=\frac{\sum\left[ (c_{1}\bigotimes c_{2})\circ\Omega\right]}{\sqrt{\sum\left[ (c_{1}\bigotimes c_{1})\circ\Omega\right]\times\sum\left[ (c_{2}\bigotimes c_{2})\circ\Omega\right]}}$$

given that GIPs are linear combinations of original traits ($c_{1}$ and $c_{2}$, in case of ${GIP}_{i}$ $c_{j}=a_{i}^{s}$),

The contribution of each GIP into the genetic basis of original traits (the genetic variance explained by GIP) was estimated as squared genetic correlation coefficient of GIP with a given trait.

#### References

1.	Deelen, J. *et al.* A meta-analysis of genome-wide association studies identifies multiple longevity genes. *Nat. Commun.* **10**, (2019).

2.	Timmers, P. R. H. J. *et al.* Genomics of 1 million parent lifespans implicates novel pathways and common diseases and distinguishes survival chances. *Elife* **8**, (2019).

3.	Zenin, A. *et al.* Identification of 12 genetic loci associated with human healthspan. *Commun. Biol.* **2**, 41 (2019).

4.	Atkins, J. *et al.* A Genome-Wide Association Study of the Frailty Index Highlights Synaptic Pathways in Aging. *A Genome-Wide Assoc. Study Frailty Index Highlights Synaptic Pathways Aging* 19007559 (2019) doi:10.1101/19007559.

5.	Williams, D. M., Jylhävä, J., Pedersen, N. L. & Hägg, S. A Frailty Index for UK Biobank Participants. *Journals Gerontol. - Ser. A Biol. Sci. Med. Sci.* **74**, 582–587 (2019).

6.	Harris, S. E. *et al.* Molecular genetic contributions to self-rated health. *Int. J. Epidemiol.* **46**, 994–1009 (2017).

7.	Millard, L. A. C., Davies, N. M., Gaunt, T. R., Davey Smith, G. & Tilling, K. Software Application Profile: PHESANT: a tool for performing automated phenome scans in UK Biobank. *Int. J. Epidemiol.* **47**, 29–35 (2018).

8.	Gibson, J. *et al.* A meta-analysis of genome-wide association studies of epigenetic age acceleration. *PLOS Genet.* **15**, e1008104 (2019).

9.	Wright, D. J. *et al.* Genetic variants associated with mosaic Y chromosome loss highlight cell cycle genes and overlap with cancer susceptibility. *Nat. Genet.* **49**, 674–679 (2017).

10.	Roberts, V., Main, B., Timpson, N. J. & Haworth, S. Genome-Wide Association Study Identifies Genetic Associations with Perceived Age. *J. Invest. Dermatol.* **140**, 2380–2385 (2020).

11.	Li, C. *et al.* Genome-wide Association Analysis in Humans Links Nucleotide Metabolism to Leukocyte Telomere Length. *Am. J. Hum. Genet.* **106**, 389–404 (2020).
