## Supplementary material for "Genetically independent phenotype analysis identifies LPA and VCAM1 as drug targets for human ageing": Description of supplementary data files

**Descriptions of Additional Supplementary Files**

**Supplementary Data 1**

**Description: Genetic and non-genetic correlations between 11 ageing-related trait GWAS.** Sample Size—Total sample size reported in the original study (cases + controls, if applicable). SNPs in panel—Number of SNPs shared between the GWAS and the High-definition likelihood inference of genetic correlations (HDL) reference panel (maximum of 1,029,876 SNPs). h^2^_SNP_—Observed scale SNP heritability estimated by HDL. SE—standard error of the estimate. Beta—Correlation estimate. Phenotypic correlation—correlation between Z scores of independent null SNP (Z < 1.96), capturing both correlations due to phenotypic similarity and sample overlap. nSNP—Number of independent null SNPs used in the phenotypic correlation calculation.

**Supplementary Data 2**

**Description: Genetic correlations of core ageing-related trait GWAS and GWAS-MAP phenotypes.** [Shared] Genetic correlations showing no evidence of heterogeneity between ageing-related traits. rg_meta—fixed-effect inverse-variance genetic correlation of all six core ageing-related trait GWAS. se_meta—standard error of the meta-analysed genetic correlation. p—p value of the genetic correlation estimate. p_het—Cochran’s Q heterogeneity P value. i2—I^2^ heterogeneity statistic. pmid—PubMed identifier. [Unique] Genetic correlations which are uniquely strong or weak for a single ageing-related trait. outlier—ageing-related trait showing the uniquely strong or weak genetic correlation with the GWAS-MAP trait, compared to the other ageing-related traits. rg_outlier—genetic correlation estimate of the outlier trait with the GWAS-MAP trait. se_outlier—standard error of the estimate. rg_meta—fixed-effect inverse-variance-weighted meta-analysis of the genetic correlation estimates of the remaining ageing-related traits (i.e. excluding the outlier). se_meta—standard error of the meta-analysed genetic correlation estimate. i2_omit—I^2^ heterogeneity statistic after excluding the specified trait, where “fathlife” refers to Father lifespan, “resilience” refers to the inverse of Frailty index, “healthspan” refers to Healthspan, “longevity” refers to Exceptional longevity, “mothlife” refers to Mother lifespan, and “selfhealth” refers to Self-reported health. [All correlations] All genetic correlation data. [All heterogeneity] All statistics on heterogeneity between ageing-related trait GWAS.

**Supplementary Data 3**

**Description: Genetic correlations of the six ageing GIPs and GWAS-MAP phenotypes.** rg—Genetic correlation estimate. se—Standard error for the estimate. p—nominal, Wald-ratio P value for the correlation estimate. p_adj—Wald-ratio P value, Bonferroni-adjusted for testing 729 traits and six GIPs. pmid—PubMed identifier.

**Supplementary Data 4**

**Description: Changes in lead GIP1 SNP effects when excluding one of the core ageing-related trait GWAS.** Nearest gene(s)— Nearest gene or cytogenetic band to the lead GIP1 SNP in the locus. rsID—lead SNP of the locus in the main GIP1 GWAS. Difference—Difference in effect size of the SNP in the main GIP1 GWAS and the GIP1 GWAS constructed from five core ageing-related trait GWAS (excluding the listed trait). SE—Standard error. P—Nominal two-sided P value for the difference. Loci with differences passing Bonferroni correction for 27 tests are marked in bold. N Het—Number of scenarios in which the SNP effect size is significantly different when excluding one of the traits.

**Supplementary Data 5**

**Description: Lookup of GIP1 loci in the GWAS catalog and PhenoScanner.** Locus—Nearest gene or cytogenetic band to the lead GIP1 SNP in the locus. Trait—Shortened trait name. rsID—lead GIP1 SNP of the locus. A1—Effect allele, associated with increased GIP1. Proxy—SNP in linkage disequilibrium (r^2^_EUR_ ≥ 0.8) with the lead SNP, for which the trait association statistics are reported. P.A1—Effect allele for the proxy, matching the lead SNP A1 allele. R2—Linkage disequilibrium r^2^ value. Beta1—Reported effect size associated with carrying one copy of the P.A1 allele. SE—Standard error. P—Reported P value. Units—Reported units for the effect size. PMID—PubMed identifier.

**Supplementary Data 6**

**Description: PASCAL gene enrichment analysis.** gene—HGNC gene symbol. entrez-id—NCBI Entrez identifier for the gene. chr—chromosome. start—Base-pair position of the start of the coding region of the gene (GRCh37). end—Base-pair position of the end of the coding region of the gene (GRCh37). n_snps—Number of SNPs within the gene region shared between the PASCAL reference and the GIP1 GWAS. p—Enrichment P value. q— Benjamini-Hochberg-adjusted P value.

**Supplementary Data 7**

**Description: PASCAL gene set enrichment analysis.**  [Hallmark] Hallmark (C1) gene sets from the Molecular Signatures Database. Pathway—Gene set name. Nominal—Chi-square-based enrichment statistics. Empirical—Permutation-based enrichment statistics. P—Uncorrected P value for enrichment. FDR—Benjamini-Hochberg-adjusted P value. [GO clusters] Gene ontology (C5.BP) pathways (FDR < 5%), clustered based on pathway Jaccard similarity. Cluster—Pathway grouping based on the minimised BIC criterion by mCLUST. [GO all] All Gene ontology (C5.BP) pathways tested by PASCAL.

**Supplementary Data 8**

**Description: Technical details and sensitivity tests of the proteome-wide Mendelian randomisation and colocalisation analysis.** [MR] Statistics for all 857 exposures. exposure—Protein used as exposure. nsnp—Number of SNPs used as instrumental variable for the exposure. Method—method used to calculate the MR effect: Wald ratio for single instrument SNPs, inverse-variance-weighted test for multiple instruments. beta—MR effect estimate. se—standard error of the MR effect estimate. p—nominal, two-sided P value for the MR effect. fdr—Benjamini-Hochberg-adjusted P value. p_het—P value for heterogeneity (requires at least two SNPs). intercept—MR-Egger intercept estimate (beta), standard error (se), and P value (p), if enough instruments were available. [LD Check] Lookup of the 30 most significant GIP1 SNPs surrounding the lead pQTL instrument. lead—the pQTL used as instrument for the exposure. rsid—SNP identifier of the 30 most significant GIP1 SNPs near the lead pQTL. chr—chromosome. pos—Base-pair position (GRCh37). p—Nominal two-sided P value of the SNP association with GIP1. r2—linkage disequilibrium r^2^ of the SNP with the pQTL. check—check whether the SNP is at least r2 > 0.8 with the pQTL. pass—does the locus pass the LD check? [Colocalisation] coloc test for the probability that the GIP1 GWAS and pQTL GWASsignals colocalise. nsnp—number of SNPs used for the coloc test. PP—Posterior probability of the hypothesis, where H0 is the hypothesis of no association in either GWAS, H1 is the hypothesis that the signal is present in the pQTL GWAS only, H2 is the hypothesis that the signal is present in the GIP1 GWAS only, and H4 is the hypothesis that the signal is shared between GWAS. coloc—Posterior probability of colocalisation (H4) of more than 80%? [MR Steiger] Steiger causal directionality test. r.exposure—SNP variance explained by the exposure. r.outcome—SNP variance explained by the outcome. steiger_pval—P value that the exposure and outcome have the correct causal direction. [Reverse MR] MR analysis of GIP1 on the exposure. nsnp—Number of instrumental variables shared between GWAS and used in the analysis. See [MR] for remaining header details.
